# Biological characterization and host-phage interaction of a novel *Pectobacterium* phage

**DOI:** 10.64898/2026.04.27.721058

**Authors:** Hao Yu, Yuyang Li, Haofang Wu, Huagui Gao, Huishan Wang, Lisheng Liao

## Abstract

Taro (*Colocasia esculenta* (L.) Schott) is an important vegetable and food crop in China, but in recent years, soft rot disease has frequently occurred during its cultivation and production. This disease damages the underground corms and petiole bases of taro, causing decay in the affected parts and emitting a foul odor, leading to wilting and lodging of the entire plant. This has resulted in significant economic losses to taro production in China, along with food safety issues and ecological problems caused by excessive pesticide use, making it urgent to find a green and efficient control method. Due to its specificity and environmental safety, phage therapy exhibits advantages that chemical pesticides cannot match, representing a promising alternative to chemical pesticides for controlling pathogenic bacteria. In the preliminary work of this study, a bacterial strain was isolated from taro soft rot in Shaoguan, Guangdong, and initially identified as *Pectobacterium colocasium* ZXC0623. Using this strain as the host bacterium, a *Pectobacterium* phage was screened and named QJphage. We analyzed its physicochemical properties and obtained its biological characteristics, including optimal titer, optimal infection latency period, optimal infection multiplicity, optimal storage solvent, and resistance to ultraviolet light, pH, and chloroform. Through homologous alignment analysis, eight tail fiber proteins encoded in the QJphage genome were predicted as putative receptor-binding proteins (RBPs). To validate this prediction, the corresponding genes were cloned downstream of the *egfp* gene via homologous recombination, and the resulting recombinant plasmids were transformed into a prokaryotic host to express EGFP-tagged tail fiber fusion proteins. Fluorescence detection and confocal laser scanning microscopy confirmed that the protein encoded by ORF04 functions as the RBP. Furthermore, lipopolysaccharide (LPS) was knocked out in the host strain *P. colocasium* ZXC0623. Both *ΔLPS1* and *ΔLPS2* mutants formed smaller plaques compared to the wild-type strain, and the *ΔLPS1* mutant additionally exhibited a significant reduction in plaque number, indicating that LPS serves as a receptor involved in QJphage adsorption. Finally, transcriptomic analysis during the latent period of infection focused on 20 genes predicted to be associated with phage-host receptor binding and anti-phage immune systems. The results revealed that pilin proteins act as potential reversible adsorption receptors for QJphage, while the host strain ZXC0623 also possesses a diverse repertoire of anti-phage defense systems. Collectively, QJphage exhibits stable physicochemical properties, a well-defined LPS-dependent infection mechanism, and a host with diverse defense systems, providing a foundation for the control of taro soft rot and future phage-related research.

**Importance:** Phage therapy has emerged as a highly effective biocontrol strategy against *Pectobacterium*, with its specificity making it particularly valuable. A critical aspect of this approach is the identification of phage receptors. The initial step in the phage life cycle involves adsorption to the bacterial host, beginning with reversible contact followed by irreversible binding between phage receptor-binding proteins and specific bacterial surface receptors. Potential receptors include glycolipids in the Gram-negative outer membrane, capsular polysaccharides, and various membrane proteins or appendages. In this study, we first characterized the physicochemical properties of the isolated QJphage. Through integrated transcriptomic and whole-genome analyses, we demonstrated that the LPS of *Pectobacterium* specifically interact with the tail fiber proteins of QJphage. This research provides the first evidence revealing the molecular mechanism of interaction between *Pectobacterium* and its phage, establishing a foundation for developing phage-based control strategies against soft rot diseases.

## Introduction

Bacteriophages (phages), viruses that infect bacteria, archaea, and other microorganisms, are ubiquitously distributed in nature, with a global population estimated to be nearly 10³¹ (1). They are ubiquitous in various environments, outnumbering bacteria by a factor of ten or more. Soil, for instance, harbors a high abundance of phages, up to 10⁹ per gram, with immense diversity, posing a significant threat to bacterial survival and proliferation (2). This rich phage resource plays a crucial role in shaping microbial communities and maintaining ecological balance (3, 4).

Phages have emerged as a promising biocontrol resource due to their unique biological properties. Utilizing phages to suppress soil-borne pathogens offers several advantages: high host specificity, enabling targeted action against pathogens with minimal environmental impact (5), rapid self-replication, leading to efficient lysis of pathogenic bacteria (6), and dynamic co-evolution with host bacteria, providing an adaptive countermeasure against pathogen evolution (7, 8), unlike the unidirectional development of antibiotic resistance (9, 10). These inherent antibacterial strengths position phages as a vital strategic reserve for controlling soil-borne diseases (11).

Recent years have witnessed rapid advancements in using phages to control bacterial diseases in agriculture. Phages have shown promising results in managing various plant diseases, including potato blackleg (12), kiwifruit canker (13), pear fire blight (14), tomato bacterial wilt (15), and Chinese cabbage soft rot (16). Several phage-based products are already registered for combating diseases like tomato bacterial canker and spot (17). For controlling *Pectobacterium,* Phage PP1 specifically lyses *Pectobacterium carotovorum* subsp. *carotovorum* (*Pcc*), rapidly reducing its pathogenicity and inhibiting soft rot and stem rot in cabbage, potato, and tomato (18). ΦPD10.3 and ΦPD23.1 were isolated from potato samples and target *Pectobacterium wasabiae* and *Dickeya solani*. Single-phage application can reduce potato tuber soft rot by approximately 80%, while a cocktail of both phages achieves a remarkable 95% reduction (19). Moreover, additional research has demonstrated that applying phages within one hour of pathogen exposure reduces soft rot symptoms by approximately 90% on potato leaves (20) while soaking infected potato seed tubers with phage suppresses soft rot and significantly enhances seed germination rates (21).

In this study, we focused on phages specifically lytic against *P. colocasium* isolated from taro. We aim to characterize the storage conditions, physicochemical properties, antibacterial activity, host range, and genomic information of these *Pectobacterium* phages, and to conduct preliminary explorations into their interaction mechanisms with host bacteria. Understanding these aspects will provide a foundation for developing effective phage-based biocontrol strategies against soft rot diseases in agricultural systems.

## Results

### Isolation, amplification, purification, and morphological characterization of QJphage

To obtain phages targeting *Pectobacterium*, soil samples were collected from taro fields affected by soft rot in Shaoguan, Guangdong Province, China. Using the host strain *P. colocasium* ZXC0623 and the double-layer agar method, a lytic phage was isolated and designated QJphage. Following three rounds of plaque purification, QJphage consistently formed circular, clear-edged plaques approximately 2 mm in diameter on the host bacterial lawn after 24 h of incubation at 28°C (Fig. 1A), indicating stable lytic activity. Transmission electron microscopy (TEM) was employed to determine the virion morphology. Negatively stained particles revealed that QJphage exhibits the characteristic morphology of the order *Caudovirales*. It possesses an icosahedral head with a diameter of 60.94 ± 1.0 nm and a long, non-contractile tail measuring 15.1 nm in width and 162.9 ± 1.0 nm in length (Fig. 1B). Based on these morphological features, QJphage is preliminarily classified as a member of the *Caudovirales*.

**FIG. 1.**
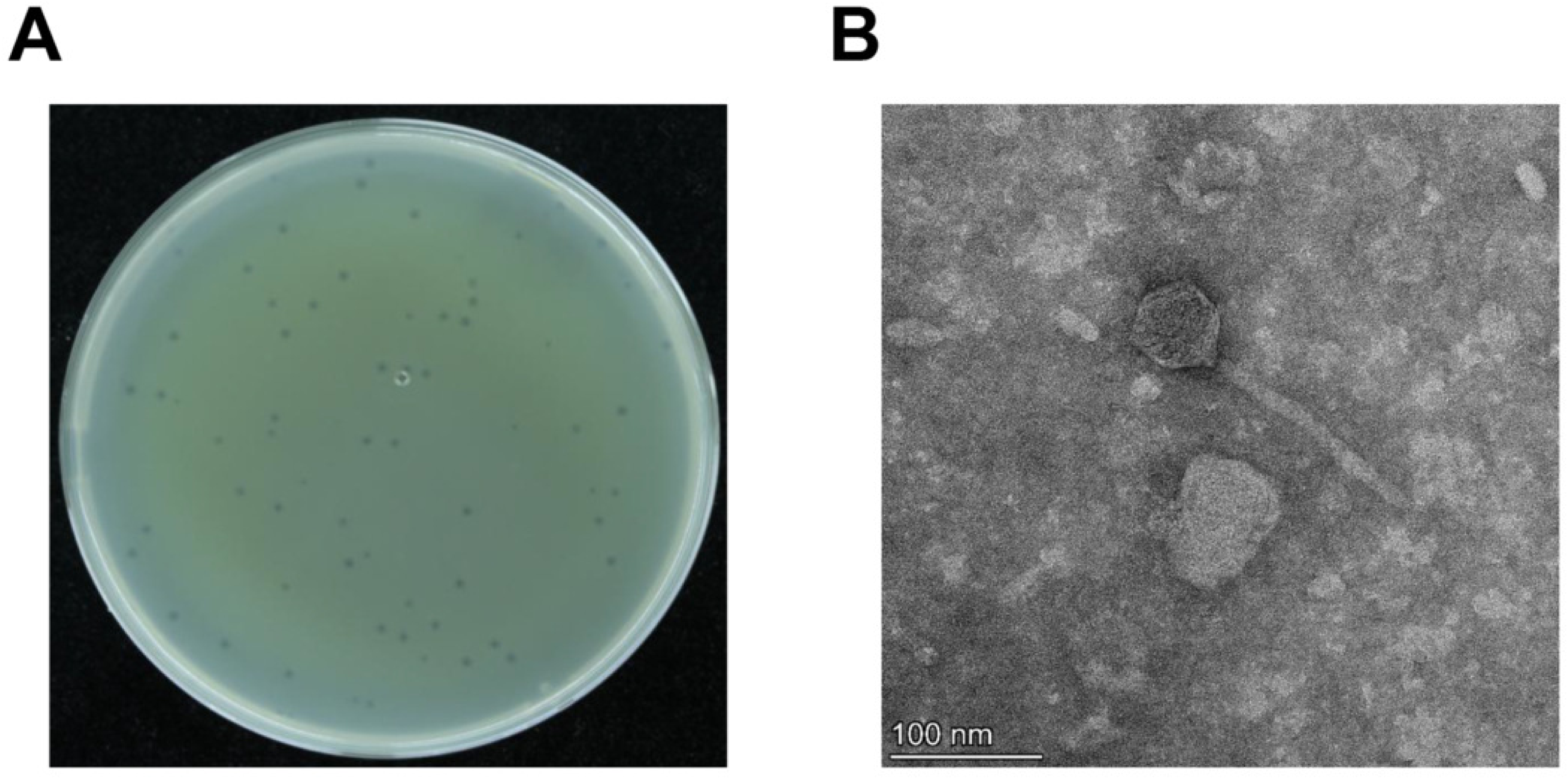
Characterization of phage biology through plaque morphology and electron microscopy. (A) Plaque morphology of the phage isolate on solid medium, exhibiting distinct and uniform lysis zones (plaque diameter: 2-3 mm) that reflect stable infectivity and host specificity. (B) Transmission electron micrographs of the phage (negative staining with 2% phosphotungstic acid; scale bar = 100 nm), revealing a typical Siphoviridae morphology with an elongated icosahedral head (∼100 nm) and a non-contractile tail (∼150 nm).

### Characterization of QJphage

To evaluate the broad-spectrum lytic activity of the phage, ten pathogenic strains isolated from taro soft rot, including six *Pectobacterium* strains (ZXC0623, LJ1, LJ2, MPC0232, MPC0225, ZXC0623a) and four *Dickeya* strains (ZXC1, CL3, ZXC054, MPC0113) (Table S1), were subjected to plaque assays. The presence or absence of lytic plaques on double-layer agar plates was used to determine the phage’s host range. QJphage exhibited lytic activity against several *Pectobacterium* strains (Table 1), including ZXC0623, MPC0225, ZXC0623a, and LJ1, but showed no detectable lysis against any of the four tested *Dickeya* strains (Table 1). These results indicate that QJphage possesses a relatively narrow host range, primarily targeting specific members of the *Pectobacterium* genus, and does not infect strains belonging to the closely related *Dickeya* genus.

**Table 1.** Host Range.

| Strains | ZXC0623 | LJ1 | LJ2 | MPC0232 | MPC0225 | ZXC0623a | ZXC1 | CL3 | ZXC054 | MPC0113 |
| --- | --- | --- | --- | --- | --- | --- | --- | --- | --- | --- |
| Host | Taro | Taro | Taro | Taro | Taro | Taro | Taro | Taro | Taro | Taro |
| QJphage | + | + | - | - | + | + | - | - | - | - |

The one-step growth curve of QJphage showed a latent period of 30 min and a rise period of 105 min (30–135 min), with a burst size of 2.95 × 10¹⁶ PFU/mL (Fig. 2A). The phage titer determined by the double-layer agar method was 4.133 × 10¹⁹ PFU/mL. The optimal multiplicity of infection (MOI) ranged from 0.1 to 1, within which QJphage produced high progeny yields (Table 2). Thermal stability assays indicated that QJphage retained full infectivity after 1 h at temperatures up to 50 °C, but was completely inactivated at 60 °C or higher (Fig. 2B). The phage also exhibited broad pH tolerance, maintaining lytic activity across pH 3.0 - 11.0 (Fig. 2C). QJphage demonstrated strong resistance to chloroform, with no significant loss of infectivity following treatment with 5% or 10% (v/v) chloroform for 10 h (Fig. 2D). Under UV exposure, the phage remained highly stable for up to 10 min, but showed a time-dependent decline in viability with longer exposure (Fig. 2E). Storage stability tests revealed that SM buffer and 30% glycerol were the most suitable preservation media for short-term storage at 4 - 28°C, whereas host bacterial suspensions and temperatures of 37°C led to substantial titer reductions (Fig. 2F). These findings characterize the key biological and physicochemical properties of QJphage, supporting its potential application as a biocontrol agent under varied environmental conditions.

**FIG. 2.**
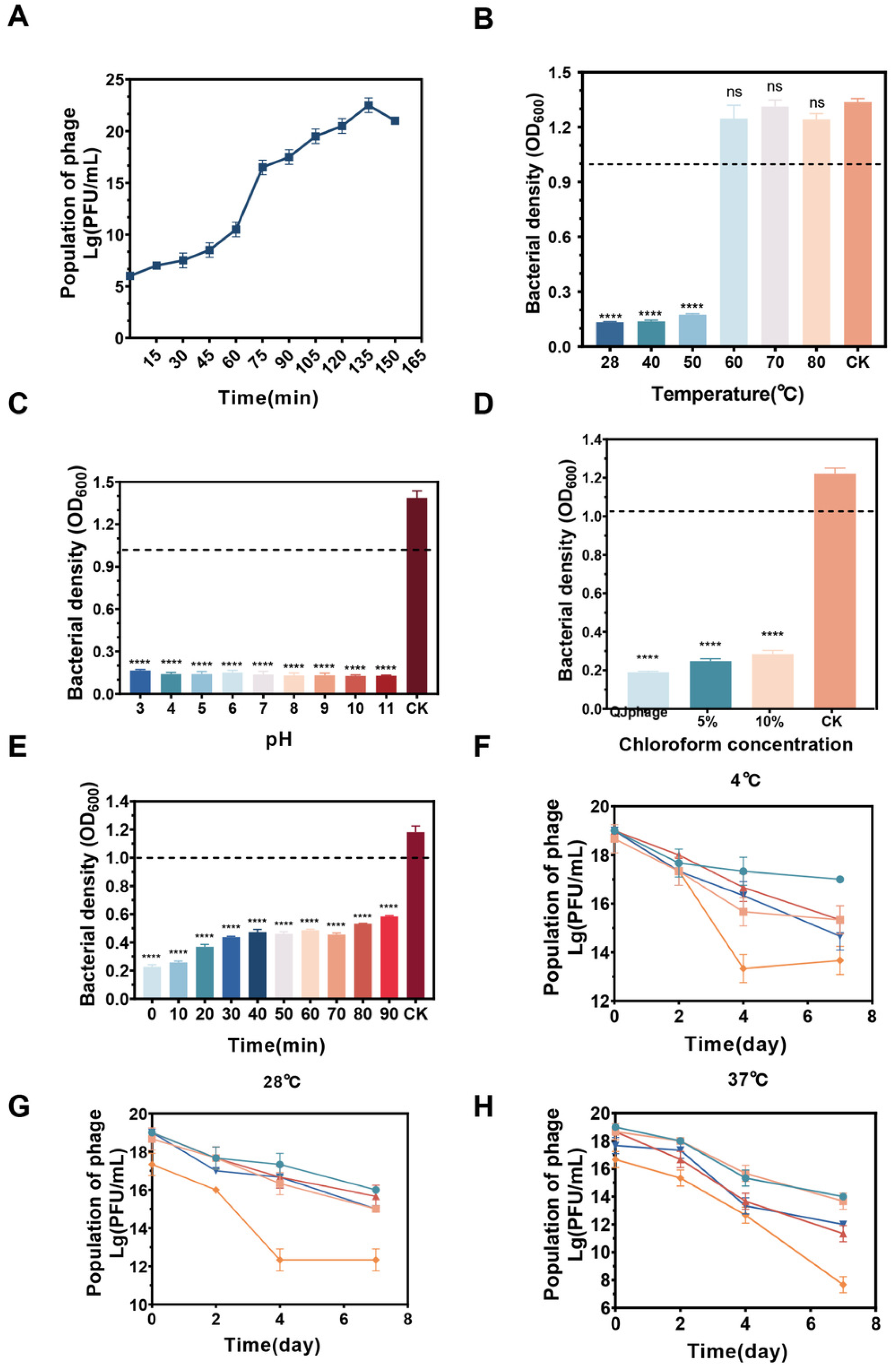
Physicochemical properties and stability of QJphage. (A) One-step growth curves of the phage in liquid culture, showing a latent phase (0–90 min) followed by rapid replication (135 min peak titer: 10⁸ PFU/mL), indicating efficient host cell lysis kinetics. (B) Thermal tolerance. QJphage was exposed to temperatures from 28°C to 80°C. Bacterial growth (OD₆₀₀) shows preserved lytic activity at `≤`50°C, with inactivation at `≥`60°C. (C) pH stability. QJphage remains stable at neutral to weakly acidic pH (5–9, lowest OD₆₀₀). Extreme pH values (`≤`4 or `≥`10) cause bacterial overgrowth. (D) Chloroform resistance. Exposure to 5% or 10% chloroform. Low-concentration chloroform (5%) has no effect, while 10% disrupts phage integrity (increased OD₆₀₀). (E) UV sensitivity. UV irradiation for `≤` 30 min shows minimal impact on bacterial density (stable OD₆₀₀). Prolonged exposure (`≥`60 min) leads to significant recovery. (F–H) Titer variation in different storage media at 4°C (F), 28°C (G), and 37°C (H). Symbols represent: circle (•), SM buffer; square (▪), 30% glycerol; upward triangle (▲), 0.85% saline solution; downward triangle ( ▾), sterile ddH₂O; diamond (♦), host bacterial suspension (ZXC0623). Statistical significance: ****, P < 0.0001; **, P < 0.01; ns, P > 0.05 (by one-way ANOVA with multiple comparisons).

**Table 2.** Determination of optimal MOI.

| MOI (PFU/CFU) | Titer (PFU/mL) |
| --- | --- |
| 100 | 4.483×10<sup>19</sup> |
| 10 | 3.025×10<sup>19</sup> |
| 1 | 2.558×10<sup>22</sup> |
| 0.1 | 2.992×10<sup>22</sup> |
| 0.01 | 1.042×10<sup>19</sup> |
| 0.001 | 1.8×10<sup>19</sup> |

### Genomic characterization and phylogenetic analysis of QJphage

The genome of QJphage was sequenced on the Illumina platform using 150 bp paired-end reads. After quality filtering, high-quality clean data were obtained. QJphage yielded 81,745,538 reads (12,201,965,132 bp), with Q20 and Q30 values of 98.50% and 94.87%, respectively, and a GC content of 47.07%. De novo assembly generated a complete linear genome of QJphage. The QJphage genome is 43,478 bp in length with a GC content of 47.07% (Table 3). Genome annotation of QJphage using PHASTEST (22) revealed a conserved and modular functional architecture. The QJphage genome comprises 58 open reading frames (ORFs), of which 42 were identified as phage genes and 16 as bacterial genes. These 42 ORFs (designated ORF01 - ORF42) were categorized into four functional modules: structural proteins, DNA packaging and processing, DNA modification and regulation, and conserved phage proteins (Fig. 3A). QJphage genome has been uploaded to the NCBI GenBank Database (GenBank accession: BankIt3069033 QJphage PZ229274).

**FIG. 3.**
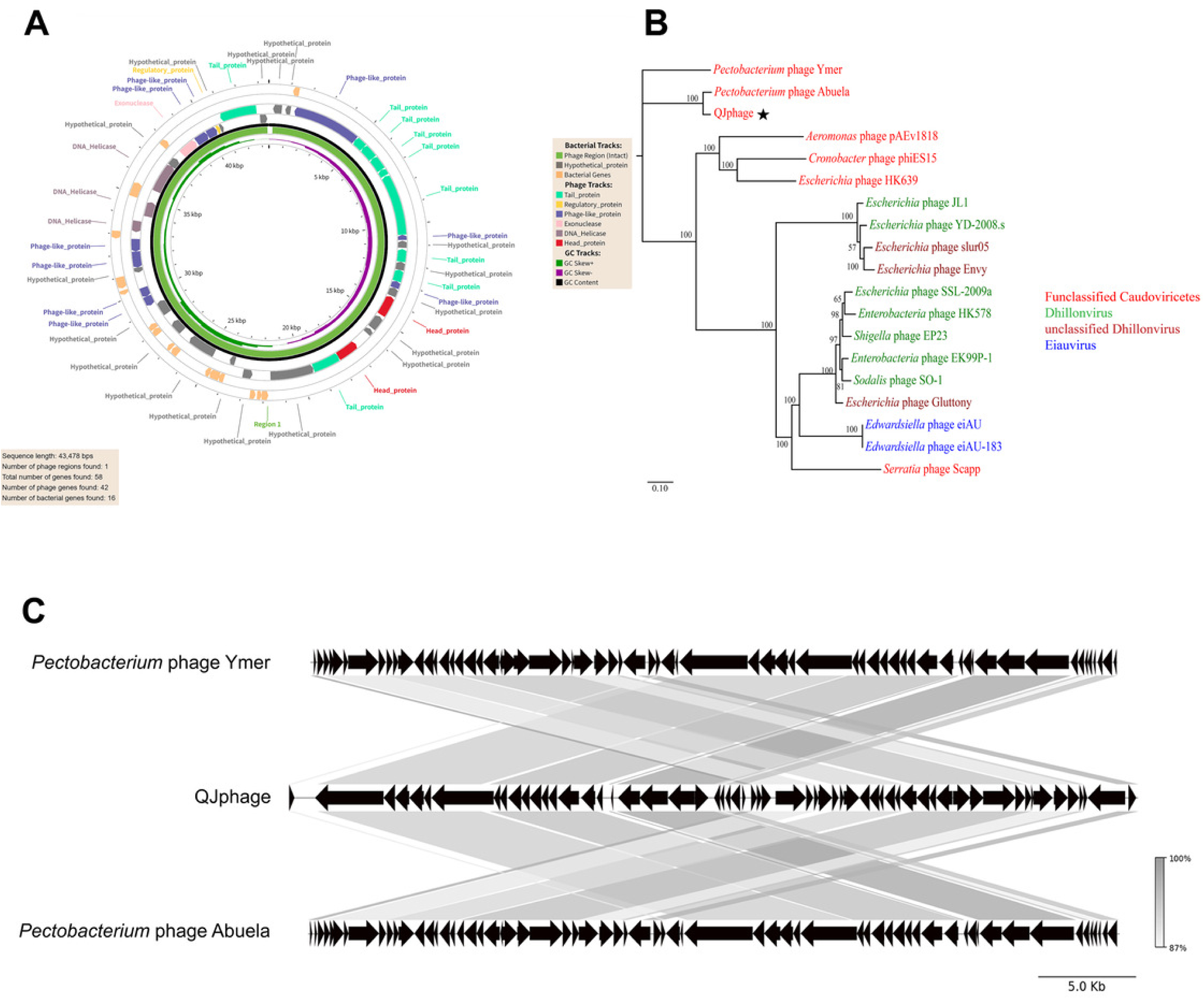
Circular genome maps and the whole-genome phylogenetic tree of QJphage. (A) Whole genome map of QJphage. (B) Branches with bootstrap support **≥**50 are labeled (node - shown values). (C) Genome synteny analysis of QJphage compared with closely related phages *Pectobacterium* phage Ymer and *Pectobacterium* phage Abuela, demonstrating a high degree of collinearity across the entire genomic region.

**Table 3.** Genomic features of QJphage.

|  |  |  |  |  |  | Illumina sequencing |  |  |  |
| --- | --- | --- | --- | --- | --- | --- | --- | --- | --- |
|  |  |  |  |  |  | Raw data |  | Clean data |  |
| Sample ID | Total | G+C | Gene | Q20 | Q30 | Total reads | Total bases(bp) | Total reads | Total bases(bp) |
|  | length(bp) | content(%) | count |  |  |  |  |  |  |
| QJphage | 43478 | 47.07 | 59 | 98.50% | 94.87% | 82376176 | 12356426400 | 81745538 | 12201965132 |

In the structural protein module, QJphage exhibits a pronounced emphasis on tail assembly machinery. ORF04, ORF05, ORF06, ORF07, ORF08, ORF11, ORF13, ORF20, and ORF41 encode tail-associated proteins that determine infection specificity through recognition of host surface receptors (23, 24). These proteins function as core sheath components involved in regulating the contraction mechanism (24, 25). Additional structural components include major head protein ORF16 and head morphogenesis protein ORF19, which contribute to capsid architecture.

In the DNA packaging and processing module, DNA replication and repair machinery to initiate genome packaging, including DNA helicases ORF33 and ORF34, a putative DNA primase/helicase ORF32, exonuclease ORF36, recombinase ORF37, and single-stranded DNA-binding protein ORF38, which protects ssDNA from degradation and enhances replication efficiency (26, 27).Regarding DNA modification and regulation, QJphage encodes a Dam family DNA N-6-adenine-methyltransferase (ORF31) that may facilitate evasion of host restriction systems through adenine methylation (28, 29). An XRE family transcriptional regulator (ORF39) is also present, potentially modulating phage gene expression during infection.A distinct group of conserved phage proteins was identified, including ORF27 (conserved phage protein), ORF28 (gp24 homolog), and ORF30 (BcepNY3gp60). These proteins, while of unknown function, are conserved across related phages and may play auxiliary roles in phage assembly or infection.Notably, no canonical holin-endolysin system was identified in the current annotation, suggesting that QJphage may employ alternative lysis mechanisms or that lysis-related genes are located within the unannotated ORFs.

Additionally, 24 ORFs encoding hypothetical proteins require further experimental validation to elucidate their biological roles.

In summary, QJphage exhibits a highly modular functional architecture. Its sophisticated tail assembly machinery and diverse DNA processing enzymes suggest a broad host range potential. These findings not only elucidate the molecular basis of phage gene functions but also provide a theoretical framework for subsequent engineering efforts, such as those aimed at biocontrol applications. Phylogenetic analysis based on the complete genome sequence placed QJphage within the informally classified Dhillonvirus group of the order *Caudovirales*, clustering closely with *Pectobacterium* phage Abuela (GenBank accession: OP748251.1) with 100% bootstrap support (Fig. 3B). Furthermore, genome synteny analysis revealed substantial genomic homology between QJphage and its close relatives, *Pectobacterium* phage Abuela and *Pectobacterium* phage Ymer (Fig. 3C). A high degree of collinearity was observed across the entire genome regions, indicating a conserved genomic architecture among these closely related phages.

### ORF04 functions as the receptor-binding protein of QJphage

To investigate the initial surface contact interaction between QJphage and its host bacterium ZXC0623, we focused on identifying receptor-binding proteins (RBPs), particularly tail fiber proteins, which mediate host adsorption. RBPs are critical determinants of host specificity and play key roles in antiviral defense and phage-based therapies. We selected 10 candidate tail fiber genes from the QJphage genome using gene annotation and PHASTEST prediction (22). In initial binding assays, EGFP-tagged tail fiber proteins were incubated with ZXC0623 (OD_600_ = 0.5). As shown in Fig. 4A, ORF04 exhibited significantly higher fluorescence intensity than other candidates, suggesting its role as the primary RBP. Concentration-gradient assays with ORF04 (0.05 - 2 mg/mL) demonstrated a dose- dependent increase in fluorescence upon incubation with ZXC0623 (Fig. 4B), supporting specific receptor interaction. This concentration-dependent binding aligns with characteristic RBP-receptor behavior.

**FIG. 4.**
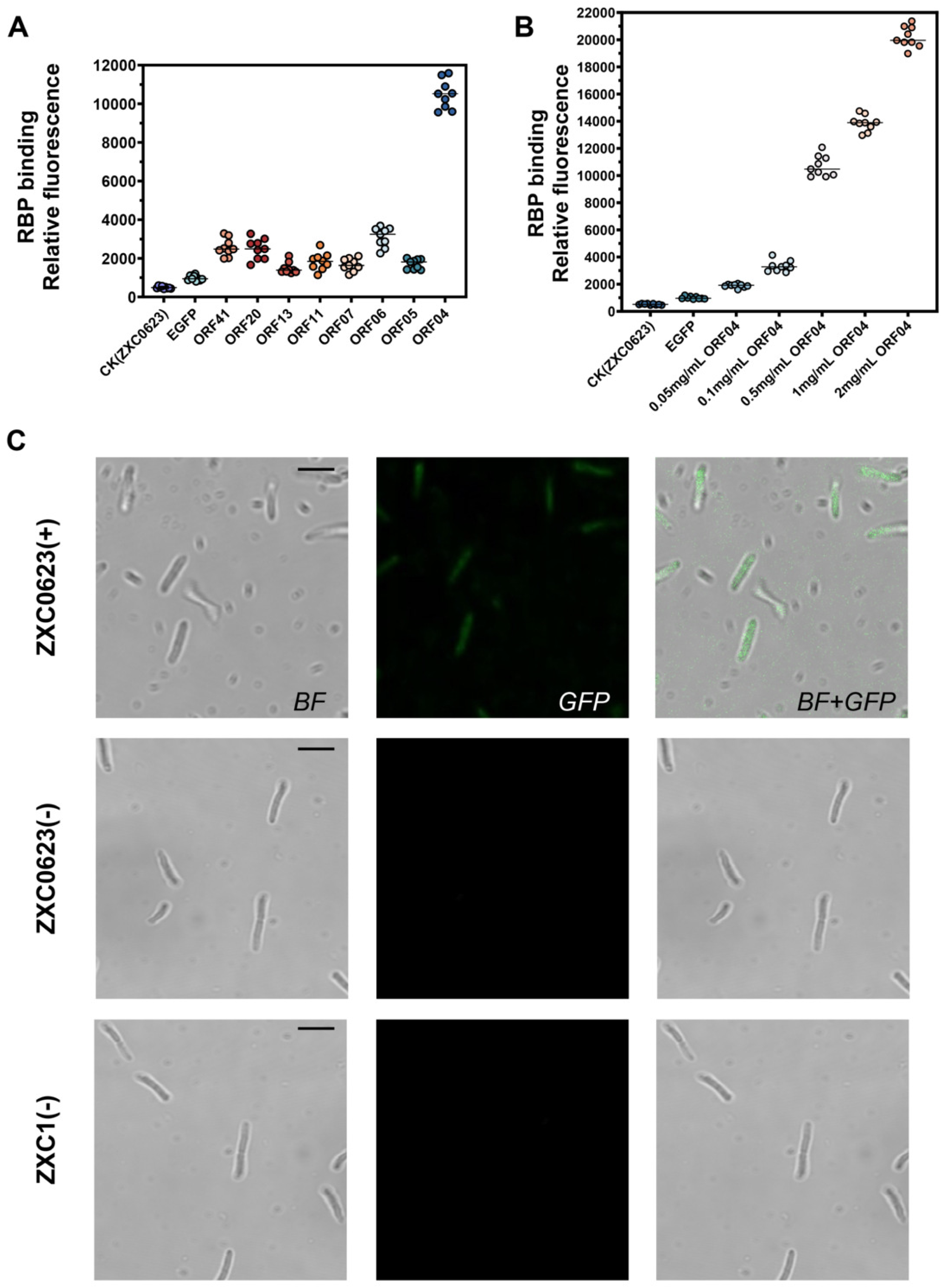
QJphage–host interaction and receptor identification on ZXC0623. (A) Fluorescence-based screening of all predicted proteins incubated with ZXC0623, followed by washing and measurement in a microplate reader, revealed strong binding affinity of the ORF04 to the host strain. (B) Concentration-dependent binding assays showed that the binding efficiency of the ORF04 to ZXC0623 increased with protein concentration. (C) Fluorescence microscopy of GFP-tagged RBP constructs incubated with wild-type ZXC0623. Strong green fluorescence was observed on bacterial cells incubated with the ORF04, while no fluorescence was detected with the empty pET28a-EGFP vector control or when the ORF04 was incubated with the non-host strain ZXC1, confirming specific binding of the ORF04 to the ZXC0623 surface.

Furthermore, the binding of the ORF04 to ZXC0623 was examined using confocal laser scanning microscopy. Approximately 90% of the wild-type ZXC0623 cells exhibited fluorescence emission at 488 nm, indicating strong binding affinity between the ORF04 and the wild-type strain (Fig. 4C). In contrast, no fluorescence was detected with EGFP protein control or when the ORF04 was incubated with the non-host strain ZXC1, confirming specific binding of the ORF04 to the ZXC0623 surface.

In addition, BLASTp reveals ORF04 homology to tail-associated proteins of enterobacterial phages, with a conserved N-terminal region and a variable C-terminus consistent with host recognition. AlphaFold2 predicts a flexible N-terminal region linked to a compact globular C-terminal domain. The protein lacks the extended β-helix typical of enzymatically active tail spike proteins, indicating it is not a depolymerase (30, 31). Its size and fold instead resemble short tail fiber or tail tip RBPs that mediate receptor binding (Fig. S3) (32).

### QJphage exhibits LPS-dependent infection

Given that lipopolysaccharide (LPS) components, including LPS and the O-antigen, commonly serve as receptor-binding targets for many Gram-negative phages (33–35), and that modifications of LPS glycan structures, which are often mediated by glycosyltransferases, have been shown to alter phage adsorption and receptor structure (36, 37), we first sought to identify candidate genes involved in LPS biosynthesis and modification in the host genome. Gene annotation and functional prediction were performed based on homology analysis against known LPS biosynthesis proteins, with particular focus on enzymes that are responsible for core oligosaccharide assembly and O-antigen ligation, as well as glycosyltransferases that may contribute to LPS structural diversification. Specifically, the *ΔLPS1* and *ΔLPS2* mutants were generated by disrupting genes encoding lipopolysaccharide heptosyltransferase (*GE03837*) and ADP-heptose-LPS heptosyltransferase (*GE03838*) (38), while the *ΔO-antigen* mutant was constructed by deleting the O-antigen ligase gene (*GE03836*). To investigate the interaction between QJphage and its host *P. colocasium* ZXC0623, we constructed mutants deficient in lipopolysaccharide (LPS) biosynthesis, including *ΔLPS1*, *ΔLPS2*, and *ΔO-antigen* strains. Plaque assays revealed that the *ΔLPS1* mutant exhibited an order-of-magnitude reduction in phage titers compared to the wild-type strain, with plaques appearing smaller and turbid. In contrast, the *ΔLPS2* mutant showed no order-of-magnitude difference but still formed smaller, turbid plaques, whereas the *ΔO-antigen* mutant exhibited no difference from the wild-type strain (Fig. 5A and 5B). These results indicate that LPS serves as a critical receptor for QJphage adsorption. To further verify that LPS is the adsorption receptor for QJphage, we incubated 0.5 mg/mL ORF04 with its host *P. colocasium* ZXC0623. Fluorescence readings showed that, compared to the wild-type strain, the *ΔLPS1* and *ΔLPS2* mutants exhibited weaker fluorescence intensity, while the *ΔO-antigen* strain showed no difference from the wild-type strain (Fig. S2).

**FIG. 5.**
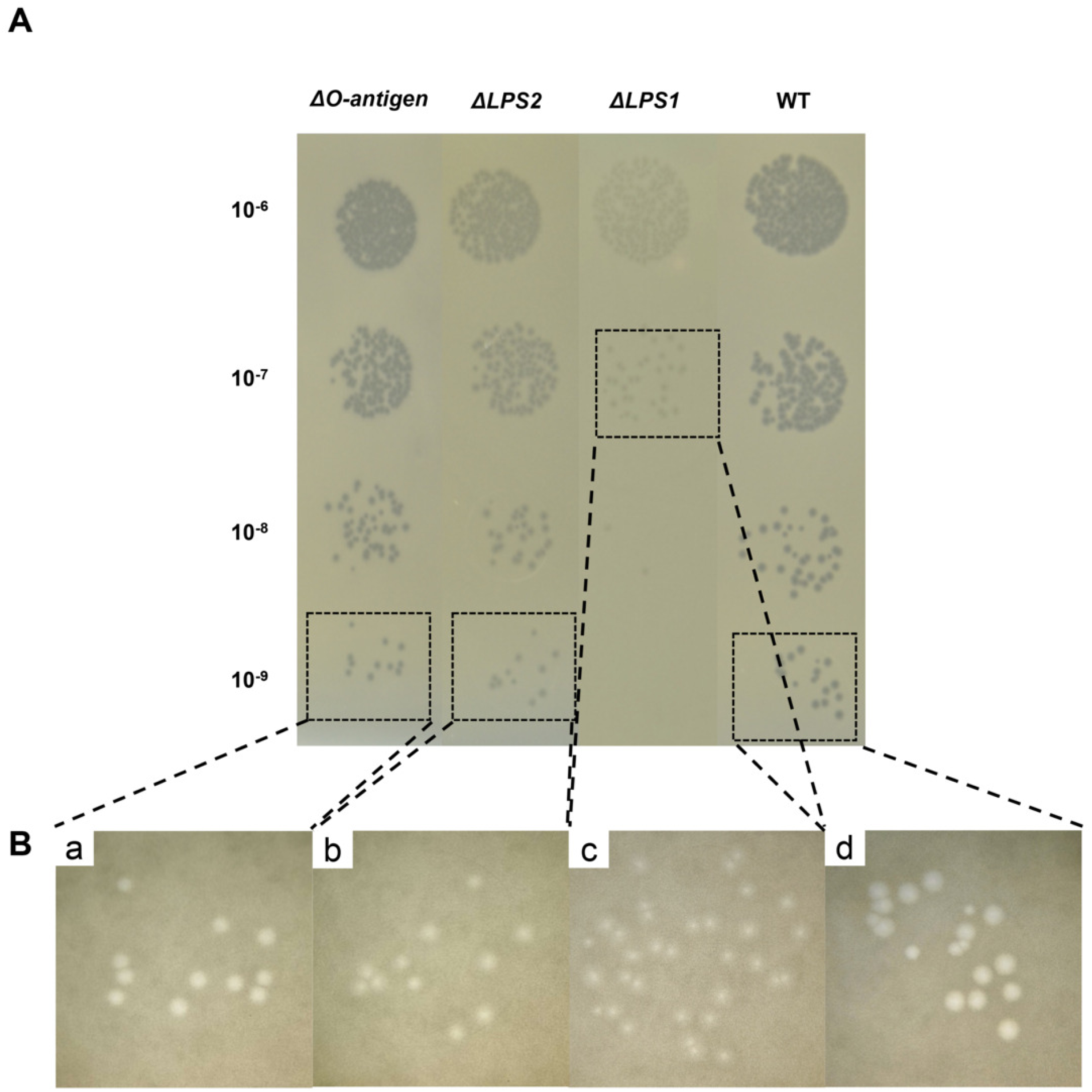
LPS serves as the adsorption receptor for QJphage. (A) Spot assays showing QJphage titers on wild-type ZXC0623 and mutant strains *ΔLPS1*, *ΔLPS2*, and *ΔO-antigen*. (B) Morphology of individual plaques formed on the mutant strains (a – c) and wild-type (**d**).

### Transcriptomic and functional analysis of *P. colocasium* ZXC0623 defense genes in response to QJphage infection

To further investigate the interaction between QJphage and *P. colocasium* ZXC0623, we performed a comparative transcriptomic analysis using ZXC0623 cells collected under QJphage stress or no stress. Differentially expressed genes (DEGs) were screened using thresholds of |log₂FC| > 1.5 and FDR < 0.05. Relative to the unstressed control, 261 DEGs were identified in the QJphage-treated group (108 significantly upregulated, 153 significantly downregulated; Fig. 6A, Table S4). KEGG pathway enrichment and GO functional annotation analysis of these 261 DEGs yielded the following results. KEGG pathway enrichment (Fig. 6B) revealed significant enrichment in metabolic pathways such as fatty acid metabolism, carbon metabolism, oxidative phosphorylation, bacterial chemotaxis, two-component systems, and fatty acid degradation. GO analysis (Fig. 6C) further indicated that DEGs were primarily enriched in functional terms including protein complex assembly, ion transport, redox processes, transmembrane transport, and associated ATPase activity, most of which are closely related to membrane structure and transmembrane transport. These findings suggest that during infection, QJphage may alter *P. colocasium* ZXC0623 physiology by disrupting host membrane function and energy metabolism. To validate the RNA-seq data, eight representative genes were selected for qRT-PCR analysis using *rpoD* as the reference gene. The results showed high consistency between qRT-PCR and RNA-seq data (Fig. 6D), confirming the reliability of the transcriptomic analysis.

**FIG. 6.**
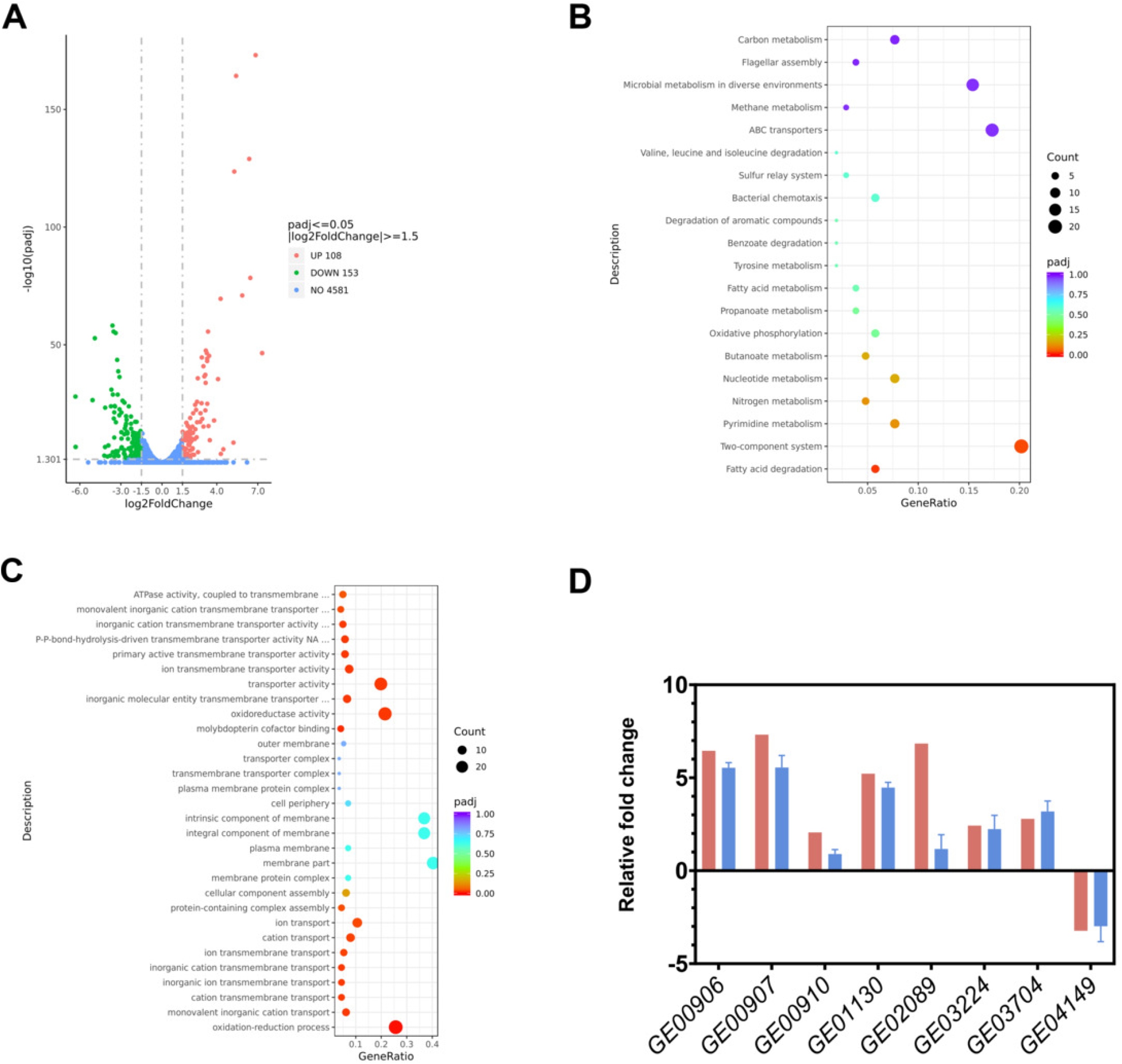
**Transcription responses and qRT**-**PCR validation.** (A) Volcano plot of DEGs in ZXC0623 (QJphage-infected vs. untreated). Genes are classified as upregulated (UP, 108), downregulated (DOWN, 153), or non-DEGs (NO, 4581), with the x-axis showing log₂ fold change and y-axis indicating −log₁₀(p-value). (B) KEGG enrichment analysis of QJphage-specific DEGs, with dot size indicating gene ratio and color representing p-value, identifying impacted metabolic pathways. (C) GO annotation dot plot of DEGs, where dot size reflects gene count and color denotes enrichment significance, highlighting biological processes perturbed by QJphage. (D) qRT-PCR validation and RNA-seq results for DEGs, showing consistent fold changes between methods. Red bars represent RNA-seq, and blue bars represent qRT-PCR.

To further elucidate the molecular response of *P. colocasium* ZXC0623 to phage infection, we focused on genes associated with membrane function, pilin component, anti-phage defense related genes (such as CBASS system), etc. Among the 261 regulated genes, 16 genes were selected and subjected to deletion and phage tolerance analysis (Table 4). Among the related knockout mutants, the pilin mutant *ΔGE02792* exhibited unstable differences in plaque formation compared to the wild-type strain (Fig. 7). Although the CBASS gene was significantly upregulated, spot assays on the CBASS-related mutants revealed no order-of-magnitude difference in phage sensitivity compared to the wild-type strain (Fig. 7). In addition, DefenseFinder (39–41) predicted 14 defense systems in the ZXC0623 genome, including the CBASS system (Table S3). The lack of a pronounced phenotype in CBASS-deficient mutants may therefore be attributed to functional redundancy or compensatory effects from other coexisting defense systems within the host.

**FIG. 7.**
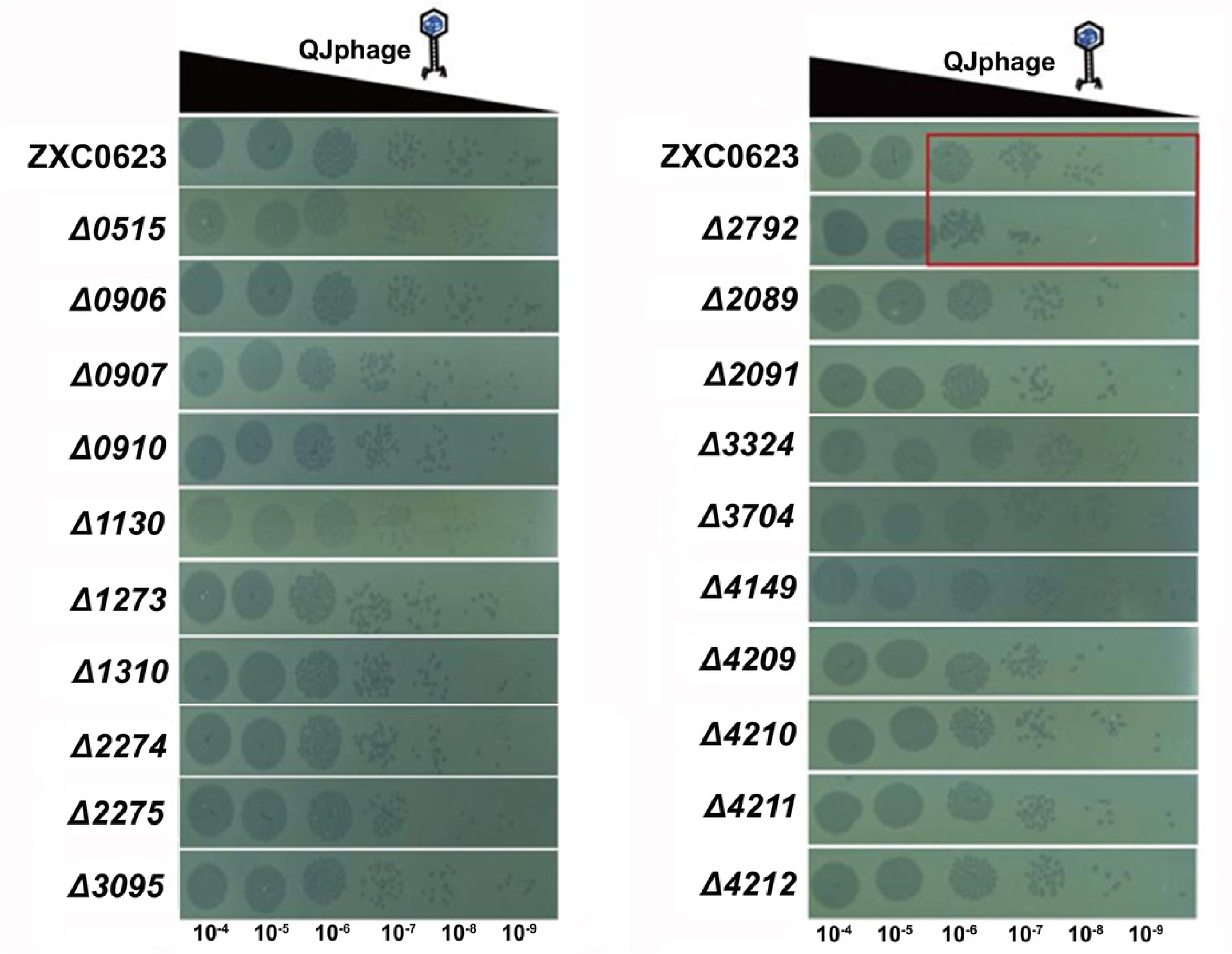
Spot assay of phage resistance in different knockout mutants. P laque assays were performed on all mutants derived from transcriptionally id entified functional genes.

**Table 4.**
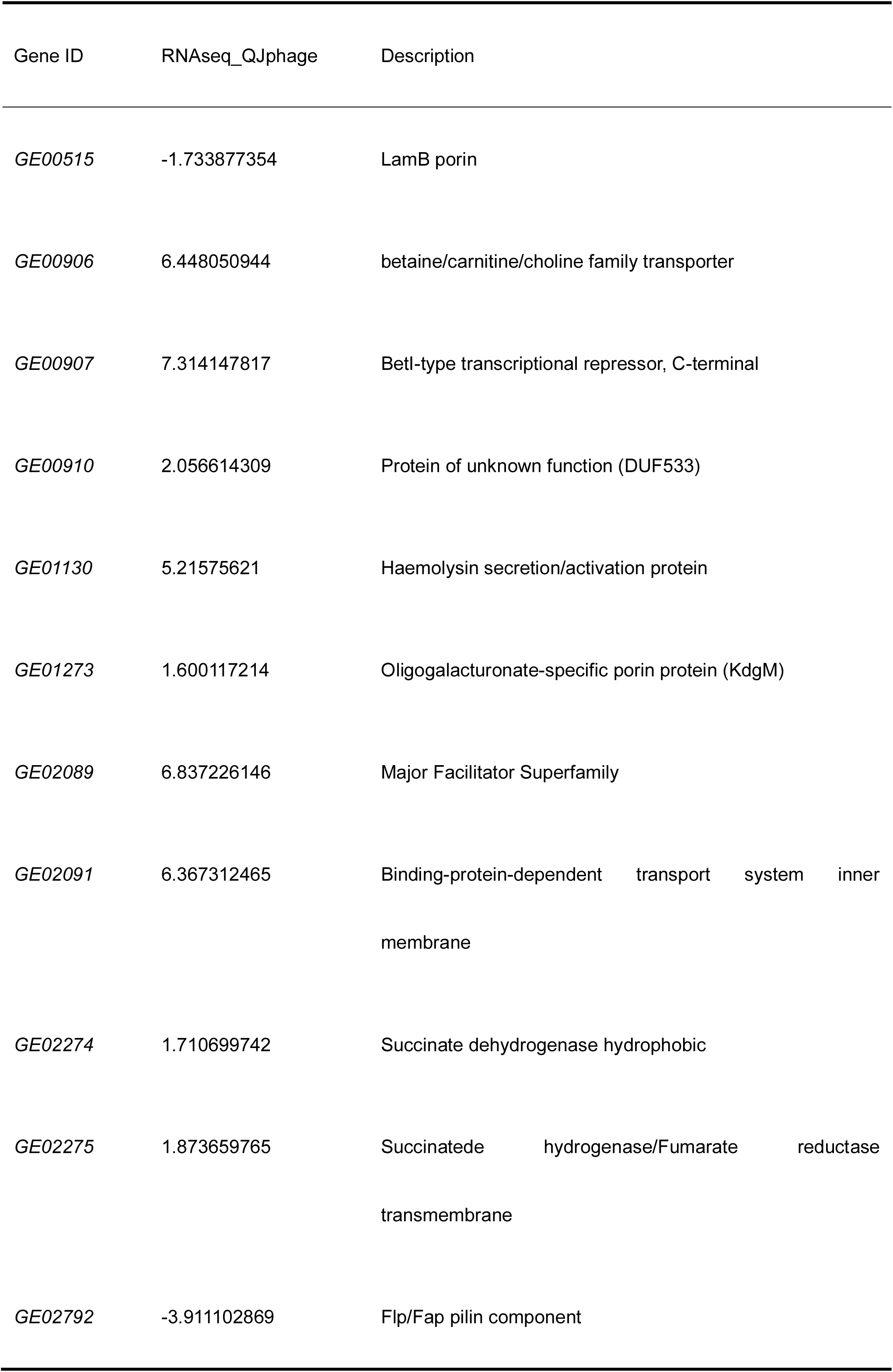
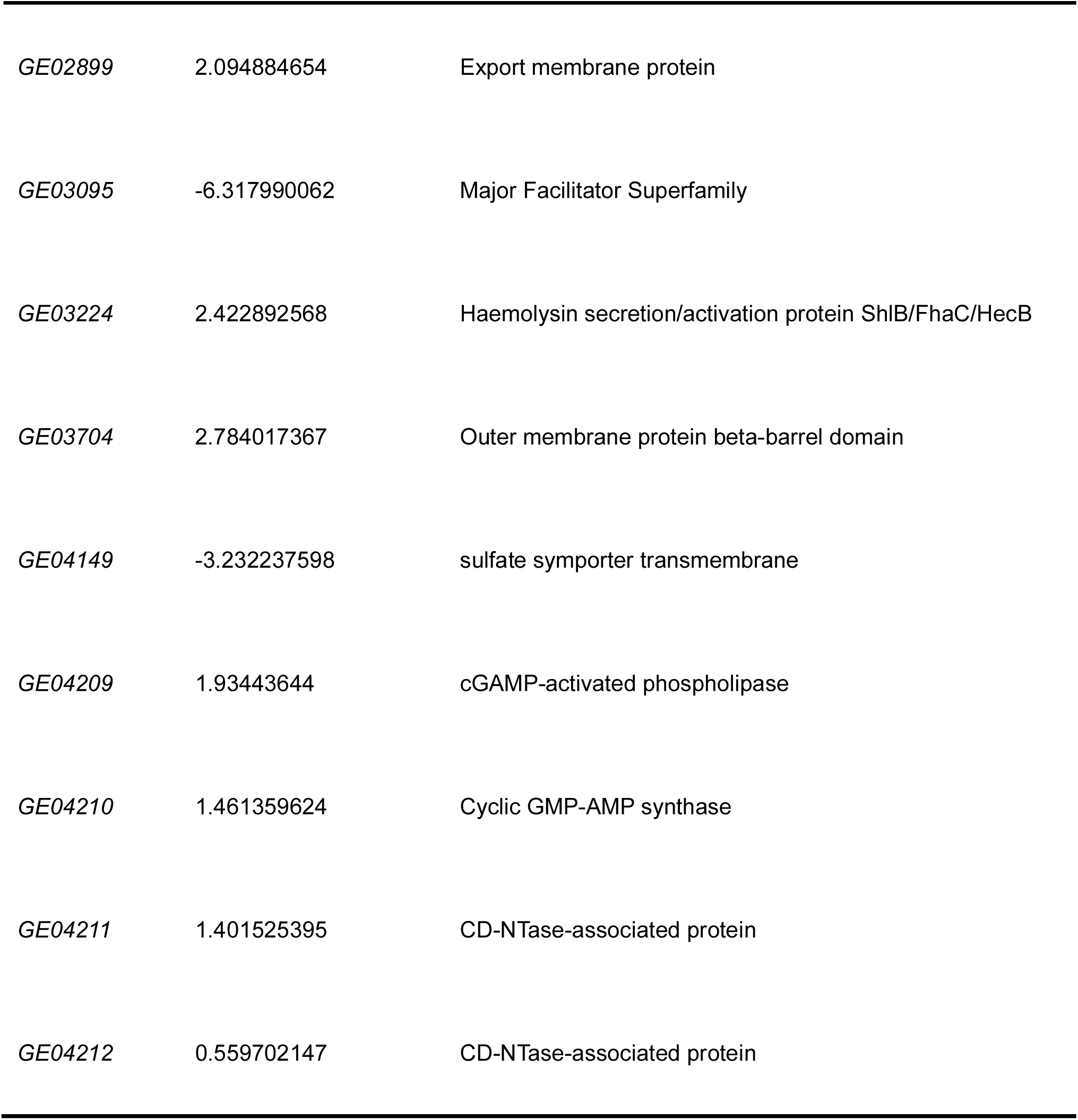
Prediction of functional genes.

| Gene ID | RNAseq_QJphage | Description |
| --- | --- | --- |
| GE00515 | -1.733877354 | LamB porin |
| GE00906 | 6.448050944 | betaine/carnitine/choline family transporter |
| GE00907 | 7.314147817 | BetI-type transcriptional repressor, C-terminal |
| GE00910 | 2.056614309 | Protein of unknown function (DUF533) |
| GE01130 | 5.21575621 | Haemolysin secretion/activation protein |
| GE01273 | 1.600117214 | Oligogalacturonate-specific porin protein (KdgM) |
| GE02089 | 6.837226146 | Major Facilitator Superfamily |
| GE02091 | 6.367312465 | Binding-protein-dependent transport system inner membrane |
| GE02274 | 1.710699742 | Succinate dehydrogenase hydrophobic |
| GE02275 | 1.873659765 | Succinate dehydrogenase/Fumarate reductase transmembrane |
| GE02792 | -3.911102869 | Flp/Fap pilin component |
| *GE02899* | 2.094884654 | Export membrane protein |
| *GE03095* | -6.317990062 | Major Facilitator Superfamily |
| *GE03224* | 2.422892568 | Haemolysin secretion/activation protein ShlB/FhaC/HecB |
| *GE03704* | 2.784017367 | Outer membrane protein beta-barrel domain |
| *GE04149* | -3.232237598 | sulfate symporter transmembrane |
| *GE04209* | 1.93443644 | cGAMP-activated phospholipase |
| *GE04210* | 1.461359624 | Cyclic GMP-AMP synthase |
| *GE04211* | 1.401525395 | CD-NTase-associated protein |
| *GE04212* | 0.559702147 | CD-NTase-associated protein |

## Discussion

Phage therapy represents a sustainable and green strategy for crop protection and production, involving the use of bacteriophages to eliminate bacterial pathogens (42). The rising prevalence of antibiotic resistance has revitalized phage research (43), however, prior to application, phages must undergo rigorous safety assessment and evaluation of their interactions with target bacteria (44). High infectivity and stability (temperature, solvents, host range) are essential prerequisites for developing phages as biocontrol agents against bacterial plant diseases. Characterization of the novel *Pectobacterium* phage QJphage demonstrated its strong thermal stability and robust tolerance to UV irradiation, pH extremes, and chloroform treatment. Combined with the potential application of inhalable phage powder (45), these properties suggest that QJphage possesses considerable survivability in field environments.

Common phage receptors in Gram-negative bacteria include LPS, outer membrane proteins, flagella, pili, saccharides, and capsules (46, 47), without which phage infection cannot occur (48). LPS, a major component of the Gram-negative outer membrane, consists of O-antigen, core polysaccharide, and lipid A (49). The O-antigen can be classified into different serotypes, and its loss results in the transition from smooth to rough LPS (50, 51). This structural diversity underlies the complexity of LPS as a phage receptor, a fact underscored by several studies. For *Pseudomonas aeruginosa*, the O-antigen of serves as a critical receptor for phage phipa9, and *ΔgtaB* disrupts O-antigen synthesis, thereby blocking phage adsorption (52). Polyvalent phage GSP004 recognizes the O-antigen of LPS via its tail fiber protein ORF208, enabling infection of both *Salmonella* and *E. coli* through distinct molecular mechanisms (53). In another case, in hypervirulent *Klebsiella pneumoniae* LPS serves as a secondary receptor that triggers phage DNA injection (54). In this study, the results indicate that LPS serves as a critical receptor for QJphage adsorption, with the O-antigen likely functioning as a primary reversible receptor that significantly influences adsorption efficiency (55–57). Notably, the mRNA transcribed from host gene *GE02792*, which encodes the piliA protein, was significantly downregulated, which may reflect an adaptive response of the bacterium under phage predation pressure, whereby active downregulation of phage receptors serves as a potential survival strategy to reduce susceptibility (58). The *ΔGE02792* mutant exhibited a one-log reduction in plaque formation compared to the wild-type ZXC0623 strain, which suggests that pili may serve as an entry portal for QJphage. To date, numerous studies have demonstrated that pili can function as phage adsorption receptors. For instance, various phages have been reported to complete adsorption and infection through specific recognition of distinct types of pili, including type IV pili and F-pili (59–62).

In the ongoing arms race between bacteria and phages, bacteria have evolved diverse anti-phage defense mechanisms to protect themselves from infection (63). At the extracellular level, bacterial strategies to resist phage invasion primarily involve receptor-mediated adaptive defenses, such as receptor modification (64), receptor blocking (65), or receptor masking. Although the CBASS system was significantly upregulated during QJphage infection, single-gene knockouts of four CBASS-related genes did not increase phage sensitivity, suggesting that multiple resistance mechanisms exist in strain ZXC0623. As expected, using DefenseFinder (39–41), we identified multiple additional defense systems (e.g., Restriction-Modification, CRISPR-Cas, AbiE, Druantia, PD-T4-5, Wadjet, and Prometheus) were identified in *P. colocasium* ZXC0623 (66–72). These systems employ distinct strategies including self versus non-self discrimination, adaptive immunity, abortive infection, and metabolic disruption, thereby forming a multilayered defense network. Although QJphage evades the CBASS system, the functional redundancy or synergistic protection conferred by these additional systems may explain the absence of observable phenotypic differences in CBASS-related mutants.

From ten predicted receptor-binding proteins (RBPs), we identified ORF04 as the functional RBP. Fluorescence-binding assays showed that ORF04 binds specifically and with high affinity to host strain ZXC0623, but not to non-host ZXC1. Unlike classical RBPs (e.g., T4, P22) that often have elongated β-structures and enzymatic activity toward polysaccharides, ORF04 appears simplified and non-enzymatic, suggesting a recognition strategy based on high-affinity binding rather than receptor degradation. This likely confers strong specificity for lipopolysaccharides on the bacterial surface (30, 73). Further structural analysis of ORF04-LPS interactions is needed to understand receptor recognition and support rational engineering for host-range modification (30, 74).

The high-affinity binding of ORF04 supports both phage therapy and the development of diagnostic tools for detecting pathogenic *Pectobacterium* in crops or soil (75). It also holds promise for targeted nanoparticle drug delivery to combat antibiotic resistance (76). Field evaluation of QJphage will be necessary to assess its environmental persistence and effects on native microbial communities in taro cultivation systems (77). Meanwhile, the strong specificity of ORF04 makes it a valuable recognition element for rapid, field-deployable pathogen monitoring (78, 79). Overall, these findings establish QJphage as a targeted biocontrol agent and inform future phage-based strategies for sustainable agriculture.

## MATERIALS AND METHODS

### Bacteria strains, growth conditions and plasmids

Bacterial strains and plasmids used in this study are listed in Table S1. Soil samples were collected from taro fields in Shaoguan City, Guangdong Province. Strain ZXC0623 was isolated from taro soft rot tissue and initially identified as *Pectobacterium* sp. by 16S rRNA gene sequencing. Its pathogenicity as the causal agent of taro soft rot disease was confirmed by fulfilling Koch’s postulates.

To further resolve its taxonomic status, average nucleotide identity (ANI) analysis was performed using FastANI (https://github.com/ParBLiSS/FastANI) to compare the complete genome of strain ZXC0623 with those of closely related type strains. The results showed that strain ZXC0623 shared 98.6% ANI with *Pectobacterium colocasium* LJ1, exceeding the species delineation threshold of 95%. Therefore, strain ZXC0623 was classified as *P. colocasium*. Phage QJphage was isolated from the same soil samples. *Pectobacterium* spp. and *Dickeya* spp. were grown at 28 °C, while *E. coli* strains were grown at 37 °C in Luria–Bertani (LB) medium overnight, unless otherwise stated. Gentamicin and kanamycin were used at final concentrations of 50 μg/mL when required for selection.

### Growth Curve Analysis

To determine the growth curve of strain ZXC0623 in LB broth, the bacterium was first streaked from -80°C stock onto an LB agar plate and incubated at 28°C for 24 h. A single colony was then inoculated into 5 mL of LB medium and cultured overnight at 28°C with shaking at 200 rpm. The overnight culture was adjusted to an OD_600_ of 1.0 and diluted 1:100 into 10 mL of fresh LB medium. Aliquots (200 µL) of this suspension were transferred into a 100-well plate for automated growth curve analysis(80). The plate was incubated at 28°C for 72 h with continuous shaking, and growth data were collected and plotted using GraphPad Prism 8.

### Phage Isolation

Soil samples (10 g) were added to sterile flasks containing 90 mL of SM buffer and incubated at 28°C with shaking at 200 rpm for 12 h. After sedimentation for 24 h, the supernatant was filtered through 0.22 μm membranes. The filtrate (5 mL) was mixed with 5 mL of logarithmic-phase host culture (ZXC0623), incubated statically for 30 min, and then shaken at 200 rpm and 28°C for 12 h. The enriched phage lysate was obtained by filtering the culture through 0.22 μm membranes. For plaque observation, 0.5 mL of the phage lysate was mixed with 0.5 mL of host culture, incubated at room temperature for 30 min, combined with 6 mL of semi-solid LB agar (cooled to approximately 50°C), and overlaid onto solidified LB agar plates. The plates were incubated at 28°C for 12 h and examined for plaque formation (81).

### Phage Purification

Individual well-isolated plaques were picked and inoculated into logarithmic-phase cultures of the host strain ZXC0623. After 30 min of static adsorption at room temperature, the mixture was incubated at 28°C with shaking at 200 rpm for 12 h to amplify the phage. The resulting lysate was filtered through a 0.22 μm membrane to obtain a crude phage stock. For plaque purification, 5 mL of a logarithmic-phase host culture was mixed with 25 mL of semi-solid LB agar (cooled to approximately 50°C) and overlaid onto solidified LB agar plates. After solidification and sectoring, the filtered phage stock was serially diluted (10^-1^ to 10^-20^), and 20 μL of each dilution was spotted onto the corresponding sector. The plates were air-dried and incubated inverted at 28°C for 12–24 h. This purification process was repeated 3–5 times to obtain homogeneous phage plaques, and the resulting purified phage suspension was stored for subsequent use (81).

### Phylogenetic Tree Construction

Whole-genome sequences were used for phylogenetic analysis. Briefly, the ViPTree server (http://www.genome.jp/viptree) was employed to generate a proteomic tree based on genome-wide sequence similarity calculated by tBLASTx, allowing virus classification. The complete genome sequence of the phage was uploaded to ViPTree with the nucleic acid type set to dsDNA and the host type set to Prokaryote. A proteomic tree was constructed, and genomes of closely related phages were selected and downloaded. Multiple sequence alignment of the query genome and the selected reference genomes was performed using MAFFT (https://mafft.cbrc.jp/). A maximum-likelihood phylogenetic tree was then generated using IQ-TREE (http://iqtree.cibiv.univie.ac.at/) with default parameters. Finally, the tree was visualized and annotated using Evolview (https://evolgenius.info//evolview-v2/#login).

### Host Range Determination

To evaluate the host range of the phage against taro soft rot pathogens, its lytic activity was tested against ten *Pectobacterium* strains isolated from diseased taro plants. Four pathogenic strains were retrieved from −80 °C storage, streaked onto solid medium, and incubated at 28 °C for 24 h (Table 1). Single colonies were inoculated into 10 mL of LB broth and cultured with shaking at 200 rpm until the logarithmic growth phase was reached. For the phage spot assay, 1 mL of each bacterial culture was mixed with 6 mL of semi-solid LB agar (cooled to approximately 45 °C) and overlaid onto solidified LB agar plates. After drying, 5 µL of phage suspension was spotted onto each bacterial lawn. The plates were incubated at 28 °C for 24 h and examined for the presence of lytic zones. All assays were performed with three independent biological replicates, each including three technical replicates.

### Plaque Assay for Phage Titer Determination

Phage titer, defined as the number of infectious virus particles per milliliter (expressed as plaque-forming units, PFU/mL), was quantified using the double-layer agar method. The purified phage suspension was subjected to serial 10-fold dilutions (ranging from 10^-1^ to 10^-20^). A 20 μL aliquot of each diluted sample was spotted onto bacterial lawns prepared in soft agar overlays. The plates were incubated at 28 °C for 24 h and examined for plaque formation. The titer was calculated based on plates showing 30–300 discrete plaques per spot, using three independent replicates (81). The phage titer (PFU/mL) was determined using the following formula: Titer (PFU/mL) = (Mean plaque count × Dilution factor) / volume plated (mL).

### MOI Determination

The multiplicity of infection (MOI) is defined as the ratio of phage particles to host bacterial cells at the onset of infection. To determine the optimal MOI, a culture of the host strain ZXC0623 in the logarithmic growth phase was mixed with phages at MOI values of 0.001, 0.01, 0.1, 1, 10, and 100 (PFU/CFU). The mixtures were incubated at room temperature for 30 min, followed by shaking at 200 rpm and 28°C for 12 h. The resulting lysates were filtered through 0.22 μm membranes. The filtrates were serially diluted, and phage titers were quantified using the double-layer agar method (82). All experiments were performed with three independent biological replicates, each including three technical replicates.

### One-Step Growth Curve Analysis

A one-step growth experiment was conducted to characterize the phage’s latent period, burst period, and burst size. Host strain ZXC0623 in the logarithmic growth phase was infected with purified phage at a MOI of 0.1. The mixture was incubated at 28 °C for 15 min to allow adsorption, followed by centrifugation at 12,000 × g for 30 s. The pellet was washed three times with LB medium to remove unadsorbed phages and resuspended in pre-warmed LB medium at 28 °C. The culture was then shaken at 200 rpm at 28 °C. Samples (100 µL) were collected at 15-min intervals for 150 min, and phage titers were immediately determined using the double-layer agar method. The one-step growth curve was plotted with incubation time as the x-axis and log10 phage titer as the y-axis (83). All experiments were performed with three independent biological replicates, each including three technical replicates.

### Thermal Stability Assay

700 μL of logarithmic-phase host culture (ZXC0623) was dispensed into each well of a sterile 48-well plate, and the initial OD_600_ was recorded. Aliquots (1 mL) of purified phage stock were incubated in a water bath at 28 °C, 40 °C, 50 °C, 60 °C, 70 °C, and 80 °C for 1 h. The heat-treated phage suspensions were added to the corresponding wells at a MOI of 0.1. The plate was incubated at 28 °C with shaking at 200 rpm for 6 h, after which the final OD_600_ was measured using a microplate reader. The difference between the final and initial OD_600_ values was used to assess the residual infectivity of the phage, with lower final OD_600_ indicating maintained lytic activity. The optimal temperature range for phage stability was determined based on these measurements. Results were presented as a bar graph with incubation temperature on the x-axis and final OD_600_ on the y-axis. All experiments were performed with three independent biological replicates, each including three technical replicates.

### UV Sensitivity Assay

700 μL of logarithmic-phase host culture (ZXC0623) was dispensed into each well of a sterile 48-well plate, and the initial OD_600_ was measured and recorded. A 15 mL aliquot of the purified phage suspension was transferred to a sterile Petri dish, forming a thin layer, and placed under a UV lamp (254 nm) in a sterilized biological safety cabinet at a fixed distance of 30 cm. Samples (1 mL) were collected at 10-minute intervals from 0 to 90 min of UV exposure. Each collected sample was immediately transferred to a sterile centrifuge tube and stored in the dark at 4 °C for 30 min to prevent photoreactivation. The UV-treated phage suspensions were added to the corresponding wells of the 48-well plate at a MOI of 0.1. The plate was then incubated at 28 °C with shaking at 200 rpm for 6 h. The final OD_600_ was measured using a microplate reader. The difference between the final and initial OD_600_ values served as an indicator of phage infectivity, with smaller increases in OD_600_ compared to the control reflecting maintained lytic activity. The maximum UV exposure time that the phages could tolerate while retaining significant infectivity was determined. Results were presented as a bar graph with UV exposure time on the x-axis and the final OD_600_ of the host culture on the y-axis. All experiments were performed with three independent biological replicates, each including three technical replicates.

### Chloroform Sensitivity Assay

First, 700 µL of logarithmic-phase host culture (ZXC0623) was dispensed into each well of a 48-well plate, and the initial OD_600_ was measured and recorded. The phage suspensions were then treated with chloroform at final concentrations of 5% and 10% (v/v) and incubated at 28°C with shaking at 200 rpm for 10 h. Two control groups were established: a bacteria-only control (CK) consisting of host culture without phage addition, and an untreated phage control containing host culture with untreated phage suspension. Following treatment, the chloroform-exposed phage suspensions were added to the corresponding wells at a MOI of 0.1. The plate was incubated at 28°C with shaking at 200 rpm for 6 h, after which the final OD_600_ was measured using a microplate reader. The difference between the final and initial OD_600_ values was used to assess the residual infectivity of the phages after chloroform exposure. Results were presented as a bar graph with chloroform concentration on the x-axis and the final OD_600_ of the host culture on the y-axis, enabling evaluation of phage stability in chloroform-containing environments. All experiments were performed with three independent biological replicates, each including three technical replicates.

### pH Sensitivity Assay

700 µL of logarithmic-phase host culture (ZXC0623) was dispensed into a 48-well plate, and the initial OD_600_ was measured. Purified phage suspensions (1 mL) were adjusted to pH values ranging from 3 to 11 using appropriate buffers and incubated at 28°C with shaking at 200 rpm for 10 h. Two controls were included: a bacteria-only blank control (CK) and a positive control with untreated phage suspension. The pH-treated phage suspensions were added to the host cultures at a MOI of 0.1. The plate was incubated at 28°C with shaking at 200 rpm for 6 h, after which the final OD_600_ was measured. The difference between the final and initial OD_600_ values was used to assess phage infectivity, with lower final OD_600_ indicating maintained lytic activity. The optimal pH range for phage stability was determined based on these measurements. Results were presented as a bar graph with pH on the x-axis and final OD_600_ on the y-axis. All experiments were performed with three independent biological replicates, each including three technical replicates.

### Effect of Storage Temperature on Phage Stability

Purified phage stocks were stored at 4°C, 28°C, and 37°C. Titers were measured at 1, 2, 4, and 7 days. Long-term storage was at -80°C. Phages maintained stable titers at 4°C but showed progressive declines at higher temperatures. -80°C storage preserved full viability. Temperature critically affects phage shelf life.

### Temperature and Solvent Effects on Phage Shelf Life

Purified phage stocks were mixed 1:1 with 30% glycerol, sterile water, 0.85% saline, or SM buffer. Short-term stability was assessed at 4°C, 28°C, and 37°C over 7 days (titers measured on days 1, 2, 4, and 7). Long-term storage was evaluated at -80°C after 6 months.

### Transcriptomic Analysis of the Interaction Between Host Bacterium ZXC0623 and QJphage

For transcriptomic sequencing, a 2 mL culture of *P. colocasium* ZXC0623 at OD_600_ = 1.0 was infected with phage at a MOI of 0.1. The mixture was incubated statically for 30 min, followed by shaking at 200 rpm and 28 °C for 15 min. Bacterial cells were subsequently collected by centrifugation. Two biological replicates were prepared under identical conditions and submitted to Novogene (Beijing, China) for transcriptome sequencing.

### qRT-PCR Validation

To validate the reliability of the transcriptomic data obtained during phage infection of *P. colocasium* ZXC0623, selected differentially expressed genes were verified by quantitative real-time PCR (qRT-PCR). Total RNA was extracted from bacterial samples and reverse-transcribed into cDNA. The cDNA template was diluted 100-fold as a working solution. Reaction mixtures were prepared on ice in a 96-well plate, with each treatment including three technical replicates and the entire experiment being independently repeated three times. Prior to qRT-PCR analysis, the plates were centrifuged to remove air bubbles. The thermal cycling protocol consisted of an initial denaturation at 95 °C for 30 s, followed by 40 cycles of 95 °C for 30 s and 60 °C for 30 s. A melt curve analysis was subsequently performed (95 °C for 15 s, 60 °C for 60 s, and 95 °C for 15 s). The resulting qRT-PCR data were analyzed for differential gene expression, and the relative expression levels of target genes were calculated using the ΔΔCt method.

### Transmission Electron Microscopy of Phage Particles

Phage particles were concentrated from 100 mL of logarithmic-phase host culture (ZXC0623) infected with 1 mL of purified phage suspension. After 30 min of static adsorption at room temperature, the culture was incubated at 28°C with shaking at 200 rpm for 12 h. DNase I and RNase A were added to final concentrations of 1 μg/mL, followed by incubation at 37°C for 30 min. Chloroform was then added to 1% (v/v) to complete cell lysis during an additional 37°C incubation for 30 min. The lysate was treated with 5.84 g NaCl per 100 mL, vortexed, and incubated on ice for 1 h. The supernatant was collected by centrifugation at 12,000 × g for 10 min. PEG8000 was added to 10% (w/v), dissolved by shaking, and the solution was incubated on ice for 1 hour. The precipitate was collected by centrifugation at 12,000 × g for 10 min and resuspended in 2 mL SM buffer. The suspension was extracted twice with equal volumes of chloroform (30 s vortexing, 5,000 × g for 10 min). For TEM observation, 10 μL of the concentrated phage solution was applied to a copper grid and allowed to adsorb for 30 min. The grid was negatively stained with 2% phosphotungstic acid (PTA) for 10 min, blotted dry with filter paper, and air-dried completely at room temperature before imaging with a field emission transmission electron microscope.

### Cloning of RBP-expressing *E.coli*

QJphage genomic DNA was extracted using the Phage DNA Isolation Kit (Norgen Biotek). The gene fragment encoding the receptor-binding protein ORF04 was amplified by PCR using the extracted DNA as template. The amplified fragment was subsequently inserted into the C-terminus of the pET28a-EGFP vector via Gibson assembly. The constructed plasmid was transformed into *E. coli* DH5α competent cells using heat shock. After plasmid purification, the construct was transformed into *E.coli* BL21(DE3) competent cells for high-levprotein expression.

### Overexpressing and purification of phage receptor-binding proteins

*E. coli* BL21(DE3) strains carrying pET28a-EGFP-RBP plasmids were cultured in LB medium at 37°C until reaching mid-log phase (OD₆₀₀ = 0.5-0.6). Protein expression was induced by adding 0.3 mM Isopropyl β-D-1-thiogalactopyranoside (IPTG), followed by overnight incubation at 16°C with shaking. Cells were harvested by centrifugation at 5,000 × g for 30 min at 4°C.The cell pellet was resuspended in 40 mL of ice-cold PBS and lysed using a pre-chilled high-pressure homogenizer. After centrifugation at 5,000 × g for 15 min at 4°C, the supernatant was incubated with Ni-NTA beads for 15 min on ice. The His-tagged proteins were purified through gravity flow chromatography, including washing with 20 mM imidazole and elution with an imidazole gradient. Protein concentration was quantified using the NanoDrop method.

### RBP Fluoreader Assay

Overnight cultures of test strains were inoculated into fresh LB medium at 1% (v/v) and grown to an OD_600_ of 0.6. The cultures were then harvested by centrifugation and concentrated to an OD_600_ of 2.0 (38). Bacterial suspensions (250 µL) were mixed with 250 µL of purified RBP (0.5 mg/mL) and incubated at 28 °C with shaking at 220 rpm for 10 min. After incubation, the cells were washed by centrifugation at 13,000 × g for 2 min, and the pellets were resuspended in 150 µL of 1× PBS. Fluorescence intensity was measured using a microplate reader (Ex: 470–15; Em: 515–20).

### Confocal laser scanning microscopy of EGFP-RBP binding

Overnight cultures of test strains were inoculated into fresh LB medium at 1% (v/v) and grown to an OD_600_ of 0.6. The cultures were then harvested by centrifugation, and the cell pellets were collected and resuspended in 100 µL of 1× PBS. The resuspended cells were incubated with 1 mL of RBP solution (0.5 mg/mL) and shaken at 220 rpm at 28 °C for 1 h. After incubation, the cells were washed three times with 1× PBS by centrifugation (9,000 × g, room temperature, 5 min per wash), and the final pellet was resuspended in 100 µL of 1× PBS. A 10 µL aliquot of the suspension was used for slide preparation. Samples were observed using a Leica Stellaris 5 confocal laser scanning microscope equipped with a 5 mW laser (488– 645 nm). Bright-field imaging was performed, and GFP fluorescence signals were acquired using 488 nm laser excitation.

### Construction of Target Gene Knockout Mutants in *P. colocasium* ZXC0623

Homologous arms (approximately 800 bp upstream and downstream of the target gene) were amplified from ZXC0623 genomic DNA using a high-fidelity polymerase. The pGEX-18 vector was digested with BamHI-HF and HindIII-HF. The digested vector and homologous arms were assembled and transformed into *E. coli* DH5α competent cells. Positive clones were selected on LB plates containing gentamicin. The recombinant plasmid was then introduced into *E. coli* S17-1 and transferred into ZXC0623 via biparental conjugation. Mutants were selected on sucrose plates.

### Phage Resistance Screening of Knockout Mutants

To evaluate the phage resistance of knockout mutants, plaque assays were performed using QJphage. Knockout mutants and wild-type ZXC0623 were streaked on LB plates and incubated at 28°C for 24 h. Single colonies were inoculated into LB and cultured overnight at 28°C with shaking at 200 rpm. The cultures were diluted 1:100 in fresh LB and grown to OD_600_ = 1.0. Then, 5 mL of bacterial culture was mixed with 25 mL of semi-solid LB (cooled to 50°C) and overlaid onto solid LB plates. QJphage was serially diluted (10^-1^ to 10^-9^) in LB, and 5 µL of each dilution was spotted onto the bacterial lawns. After drying, the plates were inverted and incubated at 28°C for 24 h. Plaque formation was observed and compared between the knockout mutants and wild-type strain.

## SUPPLEMENTAL MATERIAL

Supplemental material is available online only.

TABLE S1 Strains and plasmids used in this study.

TABLE S2 Primers used in this study.

TABLE S3 Predicted phage defense systems in the host strain *Pectobacterium colocasium* ZXC0623.

TABLE S4 Differential expression analysis results for RNA-seq pilot experiment.

FIG. S1-S3

## DATA AVAILABILITY

The genome sequence of *P. colocasium* ZXC0623 has been deposited in the GenBank database under accession number JBXGRA000000000. The transcriptome sequencing data of ZXC0623 under phage infection have been deposited in the NCBI Sequence Read Archive (SRA) under BioProject accession number PRJNA1455624, with SRA accession numbers SRR38195893– SRR38195896.

## FOUNDINGS

This study was supported by Guangdong S&T Program (2026B0202190005), and the Science and Technology Plan Project in Guangzhou (2025A04J7118).

### Authorship contribution statement

H.Y., Y.L., H.W. performed experiments. H.Y., Y.L., H.G., H.W., and L.L. designed experiments, analyzed the data, and wrote the paper.

## REFERENCES

1. Hendrix RW, Smith MC, Burns RN, Ford ME, Hatfull GF. 1999. Evolutionary relationships among diverse bacteriophages and prophages: all the world’s a phage. Proc Natl Acad Sci U S A 96:2192–2197.

2. Koskella B, Brockhurst MA. 2014. Bacteria-phage coevolution as a driver of ecological and evolutionary processes in microbial communities. FEMS Microbiol Rev 38:916–931. 10.1111/1574-6976.12072.

3. Chevallereau A, Pons BJ, van Houte S, Westra ER. 2022. Interactions between bacterial and phage communities in natural environments. Nat Rev Microbiol 20:49–62. 10.1038/s41579-021-00602-y.

4. Debray R, Conover A, Koskella B. 2025. Phages indirectly maintain tomato plant pathogen defense through regulation of the commensal microbiome. ISME communications 5:ycaf065. 10.1093/ismeco/ycaf065.

5. Deng T, Ge X, Wang J. 2026. Structures of λ-like phage a8 tail tip bound to OmpC provide insight into receptor recognition. Structure. 10.1016/j.str.2026.02.002.

6. Sippel K, Velimirov B. 2025. Phage therapy: application and related problems-a review. Life (Basel, Switzerland) 16. 10.3390/life16010057.

7. Ibrahim R, Aranjani JM. 2026. Bacterial defense mechanisms against bacteriophages: an evolutionary arms race. Arch Microbiol 208. 10.1007/s00203-026-04785-x.

8. Stern A, Sorek R. 2011. The phage-host arms race: shaping the evolution of microbes. BioEssays : news and reviews in molecular, cellular and developmental biology 33:43–51. 10.1002/bies.201000071.

9. Hasan M, Ahn J. 2022. Evolutionary dynamics between phages and bacteria as a possible approach for designing effective phage therapies against antibiotic-resistant bacteria. Antibiotics (Basel, Switzerland) 11. 10.3390/antibiotics11070915.

10. Scanlan PD, Buckling A, Hall AR. 2015. Experimental evolution and bacterial resistance: (co)evolutionary costs and trade-offs as opportunities in phage therapy research. Bacteriophage 5:e1050153.

11. Ye M, Sun M, Huang D, Zhang Z, Zhang H, Zhang S, Hu F, Jiang X, Jiao W. 2019. A review of bacteriophage therapy for pathogenic bacteria inactivation in the soil environment. Environ Int 129:488–496. 10.1016/j.envint.2019.05.062.

12. Beňo F, Horsáková I, Kmoch M, Petrzik K, Krátká G, ševčík R. 2022. Bacteriophages as a strategy to protect potato tubers against dickeya dianthicola and pectobacterium carotovorum soft rot, vol 10, p. 2369.

13. Frampton RA, Taylor C, Holguín Moreno AV, Visnovsky SB, Petty NK, Pitman AR, Fineran PC. 2014. Identification of bacteriophages for biocontrol of the kiwifruit canker phytopathogen pseudomonas syringae pv. Actinidiae. Appl Environ Microbiol 80:2216–2228. 10.1128/AEM.00062-14.

14. Gayder S, Kammerecker S, Fieseler L. 2024. Biological control of the fire blight pathogen erwinia amylovora using bacteriophages. J Plant Pathol 106:853–869. 10.1007/s42161-023-01478-y.

15. Fujiwara A, Fujisawa M, Hamasaki R, Kawasaki T, Fujie M, Yamada T. 2011. Biocontrol of ralstonia solanacearum by treatment with lytic bacteriophages. Appl Environ Microbiol 77:4155–4162. 10.1128/AEM.02847-10.

16. Lee S, Vu N, Oh E, Rahimi-Midani A, Thi T, Song Y, Hwang I, Choi T, Oh C. 2021. Biocontrol of soft rot caused by pectobacterium odoriferum with bacteriophage phiPccP-1 in kimchi cabbage. Microorganisms 9. 10.3390/microorganisms9040779.

17. Czajkowski R, Roca A, Matilla MA. 2025. Harnessing bacteriophages for sustainable crop protection in the face of climate change. Microb Biotechnol 18:e70108. 10.1111/1751-7915.70108.

18. Lim J, Jee S, Lee DH, Roh E, Jung K, Oh C, Heu S. 2013. Biocontrol of pectobacterium carotovorum subsp. Carotovorum using bacteriophage PP1. J Microbiol Biotechnol 23:1147–1153.

19. Czajkowski R, Ozymko Z, de Jager V, Siwinska J, Smolarska A, Ossowicki A, Narajczyk M, Lojkowska E. 2015. Genomic, proteomic and morphological characterization of two novel broad host lytic bacteriophages ΦPD10.3 and ΦPD23.1 infecting pectinolytic pectobacterium spp. And dickeya spp. PLoS One 10:e0119812.

20. Muturi P, Yu J, Maina AN, Kariuki S, Mwaura FB, Wei H. 2019. Bacteriophages isolated in china for the control of pectobacterium carotovorum causing potato soft rot in kenya. Virol Sin 34:287–294. 10.1007/s12250-019-00091-7.

21. Voronina MV, Bugaeva EN, Vasiliev DM, Kabanova AP, Miroshnikov KA. 2019. Characterization of pectobacterium carotovorum subsp. Carotovorum bacteriophage PP16 prospective for biocontrol of potato soft rot. Microbiology (N Y) 88:451–460.

22. Wishart DS, Han S, Saha S, Oler E, Peters H, Grant JR, Stothard P, Gautam V. 2023. PHASTEST: faster than PHASTER, better than PHAST. Nucleic Acids Res 51:W443–W450. 10.1093/nar/gkad382.

23. Wang C, Zeng J, Wang J. 2022. Structural basis of bacteriophage lambda capsid maturation. Structure:30.

24. Barbirz S, Müller JJ, Uetrecht C, Clark AJ, Seckler R. 2010. Crystal structure of escherichia coli phage HK620 tailspike: podoviral tailspike endoglycosidase modules are evolutionarily related. Mol Microbiol 69:303–316.

25. Rakhuba DV, Kolomiets EI, Dey ES, Novik GI. 2010. Bacteriophage receptors, mechanisms of phage adsorption and penetration into host cell. Polish journal of microbiology / Polskie Towarzystwo Mikrobiologów = The Polish Society of Microbiologists 59:145–155.

26. Leiman PG, Arisaka F, van Raaij MJ, Kostyuchenko VA, Aksyuk AA, Kanamaru S, Rossmann MG. 2010. Morphogenesis of the t4 tail and tail fibers. Virol J 7:355.

27. Padilla-Sanchez V, Feiss RM. 2015. Mechanisms of DNA packaging by large double-stranded DNA viruses (acknowledgement). other.

28. Maynard-Smith M, Mckelvie J, Wood R, Harmer J, Ranasinghe R, Williams C, Coomber D, Stares A, Roach P. 2011. Direct and continuous fluorescence-based measurements of pyrococcus horikoshii DNA n-6 adenine methyltransferase activity. Anal Biochem 418:204–212. 10.1016/j.ab.2011.07.023.

29. Ma B, Ma J, Liu D, Guo L, Chen H, Ding J, Liu W, Zhang H. 2016. Biochemical and structural characterization of a DNA n6-adenine methyltransferase from helicobacter pylori. ONCOTARGET 7:40965–40977. 10.18632/oncotarget.9692.

30. Steinbacher S, Miller S, Baxa U, Budisa N, Weintraub A, Seckler R, Huber R. 1997. Phage p22 tailspike protein: crystal structure of the head-binding domain at 2.3 å, fully refined structure of the endorhamnosidase at 1.56 å resolution, and the molecular basis of o-antigen recognition and cleavage11edited by k. Nagai. J Mol Biol 267:865–880. 10.1006/jmbi.1997.0922.

31. Liu S, Lei T, Tan Y, Huang X, Zhao W, Zou H, Su J, Zeng J, Zeng H. 2025. Discovery, structural characteristics and evolutionary analyses of functional domains in acinetobacter baumannii phage tail fiber/spike proteins. BMC Microbiol 25:73. 10.1186/s12866-025-03790-2.

32. Pas C, Latka A, Fieseler L, Briers Y. 2023. Phage tailspike modularity and horizontal gene transfer reveals specificity towards e. Coli o-antigen serogroups. Virol J 20:174. 10.1186/s12985-023-02138-4.

33. Kiljunen S, Datta N, Dentovskaya SV, Anisimov AP, Knirel YA, Bengoechea JA, Holst O, Skurnik M. 2011. Identification of the lipopolysaccharide core of yersinia pestis and yersinia pseudotuberculosis as the receptor for bacteriophage φa1122. J Bacteriol 193:4963–4972. 10.1128/JB.00339-11.

34. Ibrahim N, Mcalister JA, Geddes-Mcalister J, Svircev AM, Weadge JT, Anany H. 2025. Phage host interactions reveal LPS and OmpA as receptors for two erwinia amylovora phages. Sci Rep 15:36527. 10.1038/s41598-025-15724-z.

35. Leprince A, Mahillon J. 2023. Phage adsorption to gram-positive bacteria. Viruses 15. 10.3390/v15010196.

36. Mann E, Ovchinnikova OG, King JD, Whitfield C. 2015. Bacteriophage-mediated glucosylation can modify lipopolysaccharide o-antigens synthesized by an ATP-binding cassette (ABC) transporter-dependent assembly mechanism. The Journal of biological chemistry 290:25561–25570. 10.1074/jbc.M115.660803.

37. Golomidova AK, Efimov AD, Kulikov EE, Kuznetsov AS, Belalov IS, Letarov AV. 2021. O antigen restricts lysogenization of non-o157 escherichia coli strains by stx-converting bacteriophage phi24b. Sci Rep 11:3035. 10.1038/s41598-021-82422-x.

38. Krusche J, Beck C, Lehmann E, Gerlach D, Daiber E, Mayer C, Müller J, Onallah H, Würstle S, Wolz C, Peschel A. 2025. Characterization and host range prediction of staphylococcus aureus phages through receptor-binding protein analysis. Cell Rep 44:115369. 10.1016/j.celrep.2025.115369.

39. Tesson F, Hervé A, Mordret E, Touchon M, D Humières C, Cury J, Bernheim A. 2022. Systematic and quantitative view of the antiviral arsenal of prokaryotes. Nat Commun 13:2561. 10.1038/s41467-022-30269-9.

40. Néron B, Denise R, Coluzzi C, Touchon M, Rocha EPC, Abby SS. 2023. MacSyFinder v2: improved modelling and search engine to identify molecular systems in genomes. Peer Community Journal 3. 10.24072/pcjournal.250.

41. Tesson F, Planel R, Egorov AA, Georjon H, Vaysset H, Brancotte B, Néron B, Mordret E, Atkinson GC, Bernheim A, Cury J. 2024. A comprehensive resource for exploring antiphage defense: DefenseFinder webservice,wiki and databases. Peer Community Journal 4. 10.24072/pcjournal.470.

42. Jaglan A, Vashisth M, Sharma P, Verma R, Virmani N, Bera B, Vaid R, Singh R, Anand T. 2024. Phage mediated biocontrol: a promising green solution for sustainable agriculture. Indian J Microbiol 64:318–327. 10.1007/s12088-024-01204-x.

43. Strathdee SA, Hatfull GF, Mutalik VK, Schooley RT. 2023. Phage therapy: from biological mechanisms to future directions. Cell 186:17–31. 10.1016/j.cell.2022.11.017.

44. Peng S, Liu Y, Liu H, Chen L, Niu X, Liang H, Higgins P, Bai Q. 2025. Understanding phage receptor-binding protein interaction with host surface receptor: the key for phage-mediated detection and elimination of pseudomonas aeruginosa. Eur J Clin Microbiol Infect Dis 44:2883–2897. 10.1007/s10096-025-05262-x.

45. Li M, Ke W, Chang RYK, Chan H. 2025. Long-term storage stability of inhalable phage powder formulations: a four-year study. AAPS J 27:128. 10.1208/s12248-025-01112-y.

46. Bertozzi Silva J, Storms Z, Sauvageau D. 2016. Host receptors for bacteriophage adsorption. FEMS Microbiol Lett 363. 10.1093/femsle/fnw002.

47. Ge H, Hu M, Zhao G, Du Y, Xu N, Chen X, Jiao XA. 2020. The "fighting wisdom and bravery" of tailed phage and host in the process of adsorption. Microbiol Res 230:126344. 10.1016/j.micres.2019.126344.

48. Gordillo Altamirano FL, Barr JJ. 2021. Unlocking the next generation of phage therapy: the key is in the receptors. Curr Opin Biotechnol 68:115–123. 10.1016/j.copbio.2020.10.002.

49. Dongyang G, Hongyue J, Linkang W, Xinxin L, Dayue H, Junna Z, Shuang W, Pan T, Xiangmin L, Ping Q. 2022. Fitness trade-offs in phage cocktail-resistant salmonella enterica serovar enteritidis results in increased antibiotic susceptibility and reduced virulence. Microbiol Spectr 10:e02914–e02922. 10.1128/spectrum.02914-22.

50. Pérez JM, Mcgarry MA, Marolda CL, Valvano MA. 2008. Functional analysis of the large periplasmic loop of the escherichia coli k-12 WaaL o-antigen ligase. Mol Microbiol 70:1424–1440. 10.1111/j.1365-2958.2008.06490.x.

51. Guo R, Jiao Y, Li Z, Zhu S, Fei X, Geng S, Pan Z, Chen X, Li Q, Jiao X. 2017. Safety, protective immunity, and DIVA capability of a rough mutant salmonella pullorum vaccine candidate in broilers. Front Microbiol 8:547. 10.3389/fmicb.2017.00547.

52. Sui B, Li X, Wang L, Xu Y, Hou Y, Khan MSI, Li N, Tan D. 2026. Intra-strain and inter-strain heterogeneity shape phage-host interactions and phenotypic adaptation in pseudomonas aeruginosa. Appl Environ Microbiol:e0005026. 10.1128/aem.00050-26.

53. Dongyang G, Shenyu P, Yuanhang Z, Shunyuan P, Xiangyu K, Jun S, Dongbo S. 2025. Polyvalent phage GSP004 recognizes o-antigen polysaccharide receptors in salmonella and escherichia coli through tail fiber protein ORF208. J Virol 99:e00810–e00825. 10.1128/jvi.00810-25.

54. Yin M, Cao L, Fu Y, Lu Y, Yan Y, Qian L, Xiang L, Zhou T, Chen H, Li Y, Zhang L. 2025. A novel tail fiber protein triggers phage DNA ejection by recognizing lipopolysaccharides of k54 hypervirulent klebsiella pneumoniae. Microbiol Spectr 13:e0217125. 10.1128/spectrum.02171-25.

55. Yu H, Wang C, Yue J, Guo W, Molineux IJ, Liu J. 2025. Structural basis of bacteriophage ur-lambda infection initiation. Sci Adv 11:eadw7914. doi:10.1126/sciadv.adw7914.

56. Heller K, Braun V. 1979. Accelerated adsorption of bacteriophage t5 to escherichia coli f, resulting from reversible tail fiber-lipopolysaccharide binding. J Bacteriol 139:32–38.

57. Dicks LMT, Vermeulen W. 2024. Bacteriophage-host interactions and the therapeutic potential of bacteriophages. Viruses 16. 10.3390/v16030478.

58. Tran VN, Burrows LL. 2026. Steering the course: targeting and exploiting surface receptors in phage therapy. J Bacteriol 208:e0037525. 10.1128/jb.00375-25.

59. Toropova K, Stockley PG, Ranson NA. 2011. Visualising a viral RNA genome poised for release from its receptor complex. J Mol Biol 408:408–419. 10.1016/j.jmb.2011.02.040.

60. Harb L, Chamakura K, Khara P, Christie PJ, Young R, Zeng L. 2020. SsRNA phage penetration triggers detachment of the f-pilus. Proc Natl Acad Sci U S A 117:25751–25758. 10.1073/pnas.2011901117.

61. Mccutcheon JG, Peters DL, Dennis JJ. 2018. Identification and characterization of type IV pili as the cellular receptor of broad host range stenotrophomonas maltophilia bacteriophages DLP1 and DLP2. Viruses 10. 10.3390/v10060338.

62. Yang L, Zhang T, Li L, Zheng C, Tan D, Wu N, Wang M, Zhu T. 2022. Characterization of pseudomonas aeruginosa bacteriophage l5 which requires type IV pili for infection. Front Microbiol 13:907958. 10.3389/fmicb.2022.907958.

63. Zhang S, Chu M, Sun X. 2025. The arms race in bacteria-phage interaction: deciphering bacteria defense and phage anti-defense mechanisms through metagenomics. Front Microbiol Volume 16–2025.

64. Wang X, Leptihn S. 2024. Defense and anti-defense mechanisms of bacteria and bacteriophages. Journal of Zhejiang University-Science B(Biomedicine & Biotechnology) 25:181–196.

65. Meidaninikjeh S, Mohammadi P, Elikaei A. 2024. Bacteriophages and bacterial extracellular vesicles, threat or opportunity? Life Sci 350:122749. 10.1016/j.lfs.2024.122749.

66. Roberts RJ, Vincze T, Posfai J, Macelis D. 2023. REBASE: a database for DNA restriction and modification: enzymes, genes and genomes. Nucleic Acids Res 51:D629–D630. 10.1093/nar/gkac975.

67. Makarova KS, Wolf YI, Alkhnbashi OS, Costa F, Shah SA, Saunders SJ, Barrangou R, Brouns SJJ, Charpentier E, Haft DH, Horvath P, Moineau S, Mojica FJM, Terns RM, Terns MP, White MF, Yakunin AF, Garrett RA, van der Oost J, Backofen R, Koonin EV. 2015. An updated evolutionary classification of CRISPR-cas systems. Nature reviews. Microbiology 13:722–736. 10.1038/nrmicro3569.

68. Dy RL, Przybilski R, Semeijn K, Salmond GPC, Fineran PC. 2014. A widespread bacteriophage abortive infection system functions through a type IV toxin–antitoxin mechanism. Nucleic Acids Res 42:4590–4605. 10.1093/nar/gkt1419.

69. Wu Y, Garushyants SK, van den Hurk A, Aparicio-Maldonado C, Kushwaha SK, King CM, Ou Y, Todeschini TC, Clokie MRJ, Millard AD, Gençay YE, Koonin EV, Nobrega FL. 2024. Bacterial defense systems exhibit synergistic anti-phage activity. Cell Host Microbe 32:557–572. 10.1016/j.chom.2024.01.015.

70. An Q, Wang Y, Tian Z, Han J, Li J, Liao F, Yu F, Zhao H, Wen Y, Zhang H, Deng Z. 2024. Molecular and structural basis of an ATPase-nuclease dual-enzyme anti-phage defense complex. Cell Res 34:545–555. 10.1038/s41422-024-00981-w.

71. Vassallo CN, Doering CR, Littlehale ML, Teodoro GIC, Laub MT. 2022. A functional selection reveals previously undetected anti-phage defence systems in the e. Coli pangenome. Nat Microbiol 7:1568–1579. 10.1038/s41564-022-01219-4.

72. Ka D, Oh H, Park E, Kim J, Bae E. 2020. Structural and functional evidence of bacterial antiphage protection by thoeris defense system via NAD(+) degradation. Nat Commun 11:2816. 10.1038/s41467-020-16703-w.

73. Kreisberg JF, Betts SD, Haase-Pettingell C, King J. 2002. The interdigitated beta-helix domain of the p22 tailspike protein acts as a molecular clamp in trimer stabilization. Protein science : a publication of the Protein Society 11:820–830.

74. Seul A, Müller JJ, Andres D, Stettner E, Heinemann U, Seckler R. 2014. Bacteriophage p22 tailspike: structure of the complete protein and function of the interdomain linker. Acta crystallographica. Section D, Biological crystallography 70:1336–1345. 10.1107/S1399004714002685.

75. Braun P, Rupprich N, Neif D, Grass G. 2021. Enzyme-linked phage receptor binding protein assays (ELPRA) enable identification of bacillus anthracis colonies. Viruses 13. 10.3390/v13081462.

76. Dokuz S, Coksu I, Acar S, Ozbek T. 2025. A precise targeting of staphylococcus aureus with phage RBP-decorated antibiotic-loaded nanoparticles. Biotechnol J 20:2300520. 10.1002/biot.202300520.

77. Thapa Magar R, Lee SY, Song Y, Lee S, Oh C. 2024. Minimal adverse effects of exogenous phage treatment on soil bacterial communities. Appl Soil Ecol 195:105250. 10.1016/j.apsoil.2023.105250.

78. Singh A, Arutyunov D, Szymanski CM, Evoy S. 2012. Bacteriophage based probes for pathogen detection. Analyst 137:3405–3421. 10.1039/C2AN35371G.

79. Ding Y, Zhu W, Huang C, Zhang Y, Wang J, Wang X. 2023. Quantum dot-labeled phage-encoded RBP 55 as a fluorescent nanoprobe for sensitive and specific detection of salmonella in food matrices. Food Chem 428:136724. 10.1016/j.foodchem.2023.136724.

80. Lee S, Vu N, Oh E, Rahimi-Midani A, Thi T, Song Y, Hwang I, Choi T, Oh C. 2021. Biocontrol of soft rot caused by pectobacterium odoriferum with bacteriophage phiPccP-1 in kimchi cabbage. Microorganisms 9. 10.3390/microorganisms9040779.

81. Stocker B. 1960. Bacteriophages. Nature 185:276. 10.1038/185276a0.

82. Hyman P, Abedon ST. 2009. Practical methods for determining phage growth parameters, p 175–202. In Clokie MRJ, Kropinski AM (ed), Bacteriophages: Methods and Protocols, Volume 1: Isolation, Characterization, and Interactions. Humana Press, Totowa, NJ.

83. Ellis EL, Delbrück M. 1939. The growth of bacteriophage. The Journal of general physiology 22:365–384.

